# Characterising ‘the munchies’; effects of delta-9-tetrahydrocannabinol (THC) vapour inhalation on feeding patterns, satiety, and macronutrient-specific food preference in male and female rats

**DOI:** 10.1101/2022.09.22.509090

**Authors:** Catherine Hume, Samantha L. Baglot, Lucia Javorcikova, Victoria Melts, John B. Bieber, Matthew N. Hill

## Abstract

With approximately 4% of the world’s population using cannabis, there is need to fully understand how cannabis impacts our health. It is universally known that increased food intake is a side effect of cannabis use, also known as ‘the munchies’, and it has been established that delta-9-tetrahydrocannabinol (THC), the major psychoactive cannabinoid in cannabis, drives these feeding effects. These appetitive effects of cannabis or THC have been modeled in humans and rodents but have not been fully explored. Therefore, the aim of this study was to use a translational pre-clinical model to characterise how inhalation of vapour from a THC-dominant cannabis extract alters daily feeding patterns and macronutrient-specific food preferences, with focus on microstructural feeding pattern analysis and potential sex differences.

We exposed adult male and female Sprague-Dawley rats to THC-dominant cannabis vapour or vehicle vapour daily, then gave rats access to different foods (chow, high-carbohydrate, and/or high-fat food) and post-vapour feeding patterns measured. To study macronutrient-specific food preferences, rats were given a post-vapour choice between a high-carbohydrate and a high-fat food. To assess satiety, rats were given pre-vapour access to a palatable preload in which they readily consume to become satiated. For some animals, blood and brain samples were collected post-vapour to measure phytocannabinoid and metabolite levels using mass spectrometry.

We show that THC vapour inhalation increases food intake in the first hour after vapour exposure, an effect that is not dependent on food type, sex or whether animals are satiated or not. These feeding effects were a result of THC vapour decreasing feeding latency and increasing feeding bout frequency. Consistently, these acute feeding effects were compensated for through reductions in subsequent food intake, and THC vapour did not promote weight gain. THC also altered macronutrient-specific food preferences, increasing high-fat food preference in standard conditions, and increasing high-carbohydrate food preference in satiated conditions so that rats no longer significantly preferred one food over the other. Following vapour exposure, female rats had higher THC and metabolite concentrations in plasma and the hypothalamus than males and showed a stronger high-fat food preference following THC vapour compared to males when given a food choice.

Overall, this study complements and builds upon previous clinical and pre-clinical data to fully characterise the effects of THC inhalation on feeding patterns and is the first to directly examine how THC alters macronutrient-specific food preferences and assess if appetitive THC-driven sex differences exist. This research sheds light on whether cannabis use can have energy-balance effects, information which is beneficial for recreational and medical cannabis users.

## Introduction

Despite the legalization and decriminalization of cannabis in many parts of the world, our understanding of the physiological effects of regular cannabis use remains incomplete. In 2020 it was estimated that over 4% of the world’s population consumed cannabis [1]. However, use varies between countries [2], where recently it was reported that 24-34% of North Americans and 15.7% Europeans use cannabis [3, 4], with 7-11% consuming cannabis daily [3]. At present, inhalation is the most common route of cannabis use, whether from smoking or vaping [3, 5], and the accessibility, range, and potency of cannabis products is ever increasing [4, 6]. Moreover, medical cannabis use is currently prevalent in North America [7], where cannabis and associated compounds have putative therapeutic benefit for treating pain, sleep disorders, nausea, cachexia, and certain mental health conditions [7, 8]. Despite this clinical potential, regular cannabis use is also associated with increased risk for negative psychological and cognitive effects [2].

It is well established that cannabis use is associated with heightened food craving and food consumption, an effect commonly referred to in popular culture as ‘the munchies’. Delta-9-tetrahydrocannabinol (THC), the main psychoactive component in cannabis, is mainly responsible for mediating these appetitive effects [9, 10], which have been documented in both human and rodent literature.

In the clinic, oral THC or inhaled cannabis increases subjective appetitive ratings for food, including ‘wanting’, ‘liking’ and ‘desire to eat’ [9, 11, 12], and increases acute food consumption [9, 11]. Furthermore, daily cannabis users show increased sweet tastant preference ratings [13], and selectively increase their consumption of sweet snack foods like bananas, cakes, and candy bars [14]. Similarly in rodent models, injected or vaporised THC can increase motivation for chow [15–17] and increase consumption [18–20]. Moreover, in rodents, these appetitive effects can also be seen with palatable foods, where injected THC can increase palatability readouts to sugar [21, 22], and injected THC or vaporised cannabis can increase intake of sweetened high-fat food [20] or chocolate ensure drink [23] respectively. Whether THC influences macronutrient-specific food preferences has not yet been directly studied, but evidence suggests that smoking cannabis is associated with increased carbohydrate consumption in humans [14, 24]. Moreover, the feeding state of the animal does not seem to influence the ability of THC to modulate food intake, as oral THC or inhaled cannabis can increase food intake in a satiety model [10, 23, 25, 26] and injected THC potentiates fasting induced chow intake [27]. However, these effects appear to be dose dependent, where lower THC doses increase food motivation [15–17] and stimulate feeding [10, 15, 20, 27], but higher doses induce hypophagia or have no feeding effects [15-17, 20, 27], presumably though other THC-associated behavioural effects, for example catalepsy and/or hypolocomotion [28]. Also, the timeframe of cannabis or THC feeding effects varies between rodent studies, likely a result of differing dose and route of administration, but the majority of studies imply that cannabis or THC alters appetitive behaviours in the first two-hours after administration [10, 20, 23, 26].

Currently it remains unclear if cannabis can influence body weight through altering energy intake, as both animal and human studies show mixed findings [14, 18, 29–31]. These cannabis feeding effects have therapeutic potential to stimulate weight gain in patients experiencing wasting associated with anorexia nervosa, chemotherapy, and HIV [32, 33]. Alternatively, in healthy individuals, cannabis driven feeding could be maladaptive, disrupting homeostatic feeding patterns and energy balance [34]. Therefore, further research is essential to assess how cannabis alters feeding behaviours, energy balance and body weight.

Using pre-clinical models is essential to fully understand the impact of cannabis on health. But many rodent studies assessing the appetitive and metabolic consequences of cannabis use systemically administer high doses of synthetic cannabinoids, reducing their translational value. As the most common route of cannabis use is inhalation [3, 5], and with the pharmacokinetic profiles of inhaled cannabis being drastically different to injected or consumed cannabis [28, 35], there is a need to study these energy-balance effects using more translationally relevant inhalation animal models, using natural cannabis products [36]. Also, injection models can show prolonged and variable effects, and are dependent on injection quality, thereby inhalation models have added benefit for improving consistency of cannabis exposure. Furthermore, rodent literature that have used inhalation models, have not fully explored how inhaled cannabis alters daily feeding patterns, or whether sex differences in cannabis-associated appetitive behaviours exist. Moreover, the effects of THC on macronutrient-specific food preferences have not been directly investigated using a controlled food choice paradigm. Therefore, the aim of this study was to fully characterise how inhalation of vapour from a THC-dominant cannabis extract influences daily eating patterns and food choices, with focus on acute microstructural feeding pattern analysis and potential sex differences. Based on previous literature, we hypothesised that THC vapour inhalation would robustly and acutely increase food intake, regardless of satiety or food type. Moreover, we predicted that THC vapour inhalation would alter macronutrient-specific food choices, specifically increasing preference for palatable, high-carbohydrate food.

## Methods

### Animals

Adult male (350-450g) and female (220-280g) Sprague-Dawley rats (Charles River Laboratories, Quebec, Canada) were single housed and maintained on a 12hr reverse dark light cycle (lights off at 08:00) in controlled conditions (temp, 21 ± 1 °C; humidity, 38 ± 1 %) with *ad lib* access to standard rodent chow (Prolab® RMH 2500; 3.34 kcal/g) and water unless otherwise stated. Prior to experiments, rats were acclimated to the reverse light cycle for two weeks and singly housed for one week. All procedures were carried out in accordance with Canadian Council on Animal Care guidelines under protocol AC19-0024; approved by The University of Calgary Animal Care Committee.

### General Experimental Design

Each experiment used 28 rats: 14 males (n=7 vehicle and n=7 THC exposure) and 14 females (n=7 vehicle and n=7 THC exposure) matched for body weight and food intake. Daily bodyweight (g) and food intake (kcal) were measured throughout experiments.

Prior to vapour exposure, rats were habituated to the vapour administration chambers without the presence of vapour, for 15-minutes once a day over three consecutive days (days 1-3). For experiments involving special diets, rats were also acclimated to the new foods for 5 consecutive days prior to vapour exposure (days 1-5). Then, rats were exposed to THC or vehicle vapour for 15-minutes once a day over three consecutive days (days 4-6 or 6-8 depending on length of habituation).

On habituation or vapour exposure days, food was removed from cages 90-minutes prior to dark phase onset (06:30) to prevent late light phase feeding from interfering with dark onset feeding patterns. Rats were then subjected to 15-minutes of habituation or vapour exposure directly before dark phase onset (07:45-08.00), and food access given at the beginning of the dark phase (08.00). Food intake was then measured at regular intervals over a four-hour period (08:00-12:00). Rats were then given chow for the remainder of the dark phase and the following light phase, and intake of this chow measured.

### Pre-vapour Satiety Procedures

For experiments characterising the effects of THC vapour on food intake in the satiated state, food was removed 90 minutes prior to dark phase onset (06:30) and rats were given access to a palatable preload (one-part standard rodent chow, 1.25 parts water, 0.25 parts sucrose by weight [25, 37]) for two-hours at dark phase onset (08:00-10:00). Rats were habituated to this procedure for five consecutive days and readily consume this palatable preload until satiated [10, 25, 37]. For experimental days, rats were subjected to preload access (08:00-10:00), vapour exposure (10.05-10.20) and food intake measurements (10.20-14:20) for three consecutive days.

### Vapour Exposure

A validated vapour inhalation system (La Jolla Alcohol Research Inc., California, USA) was used as previously described [35, 38]. Rats were subjected to inhalation of vapour from a THC-dominant cannabis extract containing 89.7% THC, 0.3% CBD, 1.7% CBG, 1.4% CBN and 0.6% CBC (Aphria Inc., Ontario, Canada) diluted to 10% THC in polyethylene glycol vehicle. Control animals were exposed to vehicle vapour (100% polyethylene glycol). Each vapour administration session lasted for 15-minutes, where rats were placed in airtight vapour chambers with the airflow set at 1 L per minute and exposed to a 10-second puff of THC or vehicle vapour every 2-minutes. This vapour administration schedule was used as it produces blood THC levels comparable to those in humans [39] and drives a robust feeding response as seen in preliminary studies.

### Post-vapour Food Intake Measurements

Immediately following vapour exposure, rats were removed from the vapour chambers, returned to their home cages, given food access and food intake measured every 30-minutes for two-hours (t=30, 60, 90 & 120-minutes), then every 30 or 60-minutes for a further two-hours (t=150, 180, 210 & 240-minutes, or t=180 & 240-minutes, respectively). This timeframe was chosen based on previous studies [10, 20, 23, 26] and the pharmacokinetic profile of THC delivered using our vapour inhalation model [35].

### Post-vapour Special Diet Access

The type of food given was dependent on the experiment. For experiments characterising the effects of THC on chow feeding patterns, rats were given access to standard rodent chow (Labdiet® Prolab® RMH 2500; 3.34 kcal/g). For experiments characterising the effects of THC vapour on intake of macronutrient-rich foods, rats were given access to an 80% carbohydrate food (‘high-carbohydrate food’; #D19102310, Research Diets Inc., New Jersey, USA; 3.8 kcal/g) or an 80% fat food (‘high-fat food’; #D19102309, Research Diets Inc., New Jersey, USA; 6.1 kcal/g) for four-hours post-vapour exposure. The macronutrient contents of the different foods used in these experiments are outlined in Table 1.

**Table 1.** Macronutrient contents of diets used. * Mostly sucrose (75%), ** Mostly coco butter (93%).

| *Name* | *Product #* | *Supplier* | *Calories provided by:* |  |  | *Energy Density*<br>*(kcal/g)* |
| --- | --- | --- | --- | --- | --- | --- |
|  |  |  | *Protein*<br>*(%)* | *Carbohydrate*<br>*(%)* | *Fat (%)* |  |
| Standard Rodent Chow | Prolab® RMH 2500 | Labdiet® | 29 | 59 | 12 | 3.34 |
| High-Carbohydrate Food | D19102310 | Research Diets Inc. | 10 | 80* | 10 | 3.8 |
| High-Fat Food | D19102309 | Research Diets Inc. | 10 | 10 | 80** | 6.1 |

For experiments characterising the effects of THC vapour on macronutrient-specific food preferences, rats were given access to both high-carbohydrate and high-fat foods [Table 1] and intake of both foods measured simultaneously over the four-hour post-vapour exposure period. To ensure equal access to each food, the rats food hoppers were fitted with a divider down the centre to keep foods separate. To control for potential food hopper side preferences, the position of the foods was switched each day.

Following the four-hour post-vapour food measurement period, food was replaced with standard chow for the remainder of the dark phase and following light phase, and intake of this chow measured.

### Post-vapour Microstructural Feeding Pattern Analysis

Food access periods were video recorded, and behaviour manually scored within the first 60-minutes after vapour exposure to quantify how THC acutely alters feeding behaviors and patterns. The latency to eat (min), time spent eating (min), number of feeding bouts (arbitrarily defined as periods of eating separated by at least 30-seconds or more), feeding bout duration (min), interbout interval (min), ingestion rate (intake (kcal) / time spent eating (min)) and satiety ratio (time spent not eating (min) / intake (kcal)) were quantified for each rat. Satiety ratio is an index to show how satiating each unit of food is through non-eating time. i.e., the higher the satiety ratio, the more satiated animals are [40–42]. These feeding behaviour parameters are routinely quantified and used to assess the effect of a drug or manipulation on feeding patterns [40–44].

Rats who didn’t eat in the first 60-minutes after vapour exposure were given a latency and interbout interval of 60-minutes and a value of 0 for time spent eating, number of feeding bouts and feeding bout duration. Ingestion rate and satiety ratio could not be calculated for rats that did not eat.

### Post-vapour Blood and Brain Sampling for Cannabinoid & Metabolite Measurement

Male and female THC vapour exposed rats from the initial chow feeding experiment (n=7 for each sex) were exposed to THC vapour for an additional day following the completion of behavioural testing. Immediately after 15-minutes of THC vapour exposure, rats were decapitated, and trunk blood collected into EDTA coated tubes and chilled on ice. The brain was quickly removed, and the hypothalamus dissected and flash frozen on dry ice. Blood was centrifuged (10,000 RPM for 10-minutes at 4 °C), and plasma collected and frozen. Blood and brain samples were stored at –80 °C until processing. Samples were also collected from a small subset of animals exposed to vehicle vapour to serve as assay controls.

Plasma and hypothalami samples were run through tandem mass spectrometry (LC-MS/MS) at the University of Calgary Southern Alberta Mass Spectrometry facility to measure THC, CBD, and THC metabolite concentrations (11-OH-THC and THC-COOH), as previously described [35, 38]. In short, 250 μl thawed plasma and homogenised whole hypothalami were added to a 20:1 solution of acetonitrile and deuterated analyte standards for THC, CBD, 11-OH-THC and THC-COOH (at a known concentration of 10 ng/ml in 1:1 methanol and water; Cerilliant, Round Rock, TX, USA). Then samples were sonicated for 20-minutes to precipitate proteins, centrifuged (1,800 RPM for 3-4-minutes at 4 °C) to remove particulates, and the supernatant transferred to glass tubes and evaporated with nitrogen gas. During evaporation, inside tube walls were washed with 250 μl acetonitrile to resuspend adhering lipids, before the samples were left to fully evaporate. Evaporated samples were resuspended in 100 μl 1:1 methanol and water and centrifuged twice (1,500 RPM for 20-minutes at 4 °C) to remove any remaining particulates, before being stored at –80 °C until processing. Samples were ran using a Eksigent Micro LC200 coupled to an AB Sciex QTRAP 5500 mass spectrometry (AB Sciex, Ontario, Canada). Analyte concentrations were calculated in relation to corresponding analyte standard readouts, and data was normalised to sample volume or weight, and shown as ng/ml for plasma and ng/g for hypothalamus.

### Data Presentation & Statistics

All data was presented as mean ± SEM. Energy intake data was normalised to body weight (kcal/kg). To portray general effects of THC vapour exposure, summary data is presented as the mean of 3 consecutive days of vapour exposure (days 4-6 or 6-8). To show effects of THC vapour exposure across the experiment, individual daily data is shown (baseline: day 3 or 5, vapour exposure: days 4-6 or 6-8). Preference was shown as food intake as a percentage of total intake, and a preference was considered significant when intake and/or preference of one food was significantly different to the other. Body weight change was calculated across vapour exposure days (days 4-6 or 6-8) and shown as a percentage of baseline body weight (day 3 or 5).

All statistics were carried out using IBM SPSS Statistics software. P≤0.05 was deemed statistically significant. Energy intake over time was analysed using a repeated measures three-way ANOVA with between subject factors as vapour treatment (vehicle and THC) and sex (male and female), and within subject factors as time (minutes or day). Differences in intake or feeding parameters for multiple foods was analysed using a repeated measures three-way ANOVA with between subject factors as vapour treatment (vehicle and THC) and sex (male and female), and within subject factors as type of food (high-fat and high-carbohydrate food). Mean body weight change and intake and feeding behaviour parameters for single food access were analysed using a two-way ANOVA with between subject factors as vapour treatment (vehicle and THC) and sex (male and female). THC and metabolite concentrations were analysed using a repeated measures two-way ANOVA with between subject factors as sex (male and female), and within subject factors as sample type (plasma and hypothalamus). Preload intake between males and females was analysed using a t-test, and effect of preload on subsequent food intake was analysed using a two-way ANOVA with between subject factors as preload treatment (preload and no preload) and sex (male and female). For repeated measures analysis, data was run through Mauchly’s test of sphericity, and if assumption of sphericity was not met, a Greenhouse-Geisser adjustment was applied. Bonferroni posthoc tests were conducted on factors containing more than 2 groups (i.e., time, day, or food type).

In the THC and metabolite data set, two male and one hypothalamus THC data points, one female plasma 11-OH-THC data point, and one male hypothalamus THC-COOH data point were excluded due to issues with sample processing. In the satiated food preference experiment, one rat in each the male and female vehicle group were excluded based on meeting outlier criteria (>2.5 standard deviations from the mean).

## Results

### THC vapour acutely increases chow intake through increasing feeding bout frequency

To assess the impact of THC vapour on chow feeding patterns in standard conditions, rats were exposed to THC or vehicle vapour and subsequent chow feeding measured.

Following vapour exposure, THC significantly increased chow intake within the first 60-minutes (interaction between time and treatment: F(7,168)=13.41, p<0.001), an effect which was irrespective of sex and consistent between vapour exposure days (no effect between days: F(2,48)=1.22, p=0.31) [Figure 1A-C]. Within this timeframe, THC altered feeding patterns [Figure 1D & Table 2] through decreasing the latency to eat (effect of treatment: F(1,24)=13.91, p=0.001) and increasing the number of feeding bouts (effect of treatment: F(1,24)=48.78, p<0.0001) to increase overall time spent eating (effect of treatment: F(1,24)=44.51, p<0.0001) [Table 2]. THC also reduced satiety ratio (effect of treatment: F(1,24)=9.82, p=0.005) [Table 2]. Other behavioural parameters remained unchanged after THC exposure, including feeding bout duration (no effect of treatment: F(1,24)=0.47, p=0.50) and ingestion rate (no effect of treatment: F(1,24)=0.65, p=0.43) [Table 2]. Furthermore, a sex difference in the amount of chow eaten and latency to eat was detected, where females generally eat less than males (effect of sex: F(1 24)=5.1, p=0.03) and have a longer latency to eat (effect of sex: F(1,24)=6.12, p=0.02) [Table 2].

**Figure 1.**
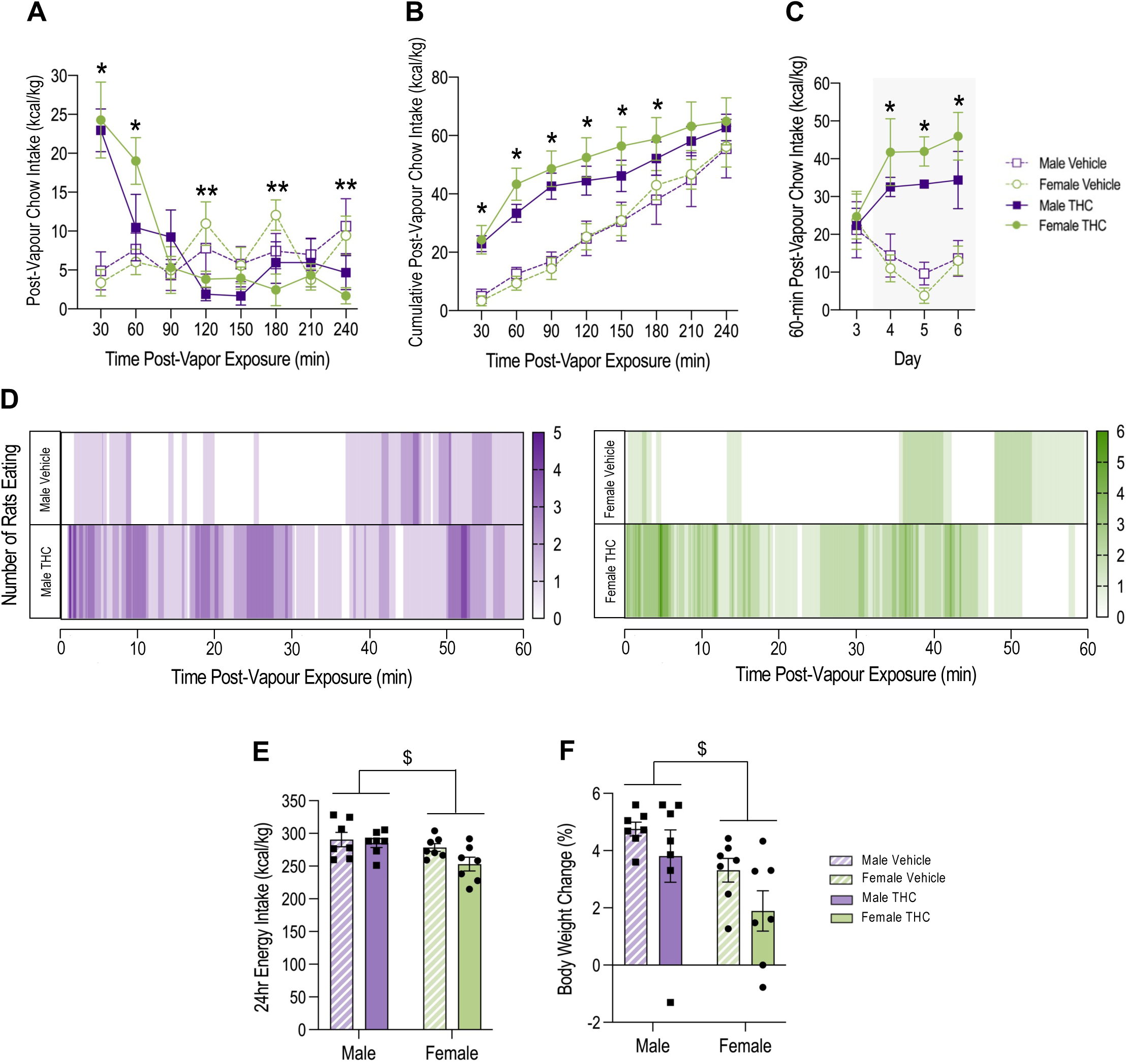
THC increases chow intake in the first 60-minutes following vapour exposure. **(A)** Day 4-6 mean post-vapour chow intake over time for THC and vehicle groups (RM 3-way ANOVA; interaction effect of time and treatment: F(7,168)=13.41, p<0.001 (post-hoc: * THC > vehicle at t=30 & 60-minutes, ** THC < vehicle at t=120, 180 & 240-minutes, p<0.05)). **(B)** Day 4-6 mean post-vapour cumulative chow intake for THC and vehicle groups (RM 3-way ANOVA; interaction effect of time and treatment: F(1.85,44.5)=4.96, p=0.013 (post-hoc: * THC > vehicle at t=30, 60, 90, 120, 150 & 180-minutes)). **(C)** Mean daily 60-minute post-vapour chow intake (day 3: baseline - no vapour exposure, days 4-6: vapour exposure - highlighted in grey) for THC and vehicle groups (RM 3-way ANOVA; interaction effect of day and treatment: F(3,72)=8.33, p<0.001 (post-hoc: * THC > vehicle on days 4-6, p<0.05)). **(D)** Representative heat maps showing the number and pattern of rats eating chow within the first 60-minutes after vapour exposure for male and female, THC and vehicle groups. **(E)** Day 4-6 mean 24-hour energy intake for THC and vehicle groups (2-way ANOVA; effect of sex: F(1,24)=6.31, p=0.019 ($ male > female)). **(F)** Body weight change (%) across vapour exposure days (day 4-6) (2-way ANOVA; effect of sex: F(1,24)=7.24, p=0.013 ($ male > female)). All intake data normalized to body weight and shown as mean ± SEM. N=7 in each group. *: effect of treatment, $: effect of sex.

**Table 2.** THC acutely increases chow intake by decreasing latency to eat and increasing feeding bout frequency to increase overall time spent eating. Day 4-6 mean feeding behaviour parameters measured within the first 60-minutes after vapour exposure. Data shown as mean ± SEM and analysed using a two-way ANOVA (for parameters with no effect, only the F and P values for treatment comparison are shown). N=7 in each group. Feeding bouts: arbitrarily defined periods of eating separated by 30-seconds or more. Ingestion rate: chow calories consumed / time spent eating. Satiety ratio: time spent not eating / chow calories consumed.

| Measure | MALE |  | FEMALE |  | Effect | F value | P value |
| --- | --- | --- | --- | --- | --- | --- | --- |
|  | Vehicle | THC | Vehicle | THC |  |  |  |
| Chow Intake (kcal) | 5.0 ± 0.9 | 13.4 ± 1.2 | 2.4 ± 0.6 | 11.3 ± 1.4 | Treatment:<br>THC > Vehicle | F(1,24)=66.61 | <0.0001 |
|  |  |  |  |  | Sex:<br>Male > Female | F(1,24)=5.1 | 0.33 |
| Time Spent Eating (min) | 5.7 ± 1.1 | 13.7 ± 1.0 | 3.2 ± 0.8 | 13.6 ± 2.2 | Treatment:<br>THC > Vehicle | F(1,24)=44.51 | <0.0001 |
| Latency to Eat (min) | 24.6 ± 4.4 | 7.6 ± 3.9 | 38.0 ± 6.5 | 16.3 ± 4.2 | Treatment:<br>Vehicle > THC | F(1,24)=13.91 | 0.001 |
|  |  |  |  |  | Sex:<br>Female > Male | F(1,24)=6.12 | 0.02 |
| Number of Feeding Bouts | 2.0 ± 0.4 | 4.9 ± 0.6 | 1.1 ± 0.4 | 5.9 ± 0.7 | Treatment:<br>THC > Vehicle | F(1,24)=48.78 | <0.0001 |
| Feeding Bout Duration (min) | 2.7 ± 0.7 | 3.3 ± 0.5 | 2.3 ± 0.6 | 2.5 ± 0.4 | No effect of<br>treatment/sex | F(1,24)=0.40 | 0.5326 |
| First Feeding Bout Duration (min) | 1.9 ± 0.7 | 1.2 ± 0.4 | 2.2 ± 0.5 | 2.1 ± 0.6 | No effect of<br>treatment/sex | F(1,24)=0.47 | 0.5005 |
| Interbout Interval (min) | 27.4 ± 3.7 | 9.2 ± 1.2 | 36.1 ± 5.4 | 10.9 ± 3.0 | Treatment:<br>Vehicle > THC | F(1,24)=30.76 | <0.0001 |
| Ingestion Rate (kcal/min) | 1 ± 0.1 | 1 ± 0.1 | 0.8 ± 0.1 | 0.9 ± 0.1 | No effect of<br>treatment/sex | F(1,24)=0.65 | 0.428 |
| Satiety Ratio (min/kcal) | 24.2 ± 14.9 | 3.5 ± 0.6 | 24.7 ± 8.8 | 4.6 ± 1.0 | Treatment:<br>Vehicle > THC | F(1,24)=9.82 | 0.0045 |

This acute THC-driven increase in chow feeding was compensated for through a subsequent reduction in chow intake so that there was no difference in the total amount of chow eaten in the 240-minute period following vapour exposure between vehicle and THC groups (no effect of treatment: F(1,24)=1.08, p=0.31) [Figure 1B]. Consequently, THC vapour exposure had no significant effect on total 24-hour energy intake (no effect of treatment: F(1,24)=3.06, p=0.09) [Figure 1E] or body weight change across vapour exposure days (no effect of treatment: F(1,24)=3.59, p=0.07) [Figure 1F]. Unrelated to treatment, a general sex difference was detected, where females consumed less energy daily (effect of sex: F(1,24)=6.31, p=0.019) [Figure 1E] and gained less body weight (effect of sex: F(1,24)=7.24, p=0.013) [Figure 1F] compared to males.

### Females have higher THC and metabolite concentrations in the blood and brain compared to males

To verify treatment exposure, rats were exposed to vapour and blood and brain samples collected to measure cannabinoid and metabolite levels.

In vehicle samples, concentrations of THC, CBD, 11-OH-THC and THC-COOH were undetectable. Following THC vapour exposure, THC, 11-OH-THC and THC-COOH were detected in moderate concentrations in the plasma and hypothalamus [Figure 2]. Compared to males, females had higher plasma and hypothalamus concentrations of THC (effect of sex: F(1,9)=15.19, p=0.004) [Figure 2A], 11-OH-THC (interaction of sex and sample: F(1,11)=8.58, p=0.014) [Figure 2B], and THC-COOH (interaction of sex and sample: F(1,11)=7.79, p=0.018) [Figure 2C]. In males and females, the hypothalamus showed higher concentrations of THC (effect of sample: F(1,9)=10.62, p=0.01) [Figure 2A] and 11-OH-THC (interaction of sex and sample: F(1,11)=8.58, p=0.014) [Figure 2B], and higher concentrations of THC-COOH in females only (interaction of sex and sample: F(1,11)=7.79, p=0.018) [Figure 2C]. Regardless of sample or sex, CBD concentrations were undetectable.

**Figure 2.**
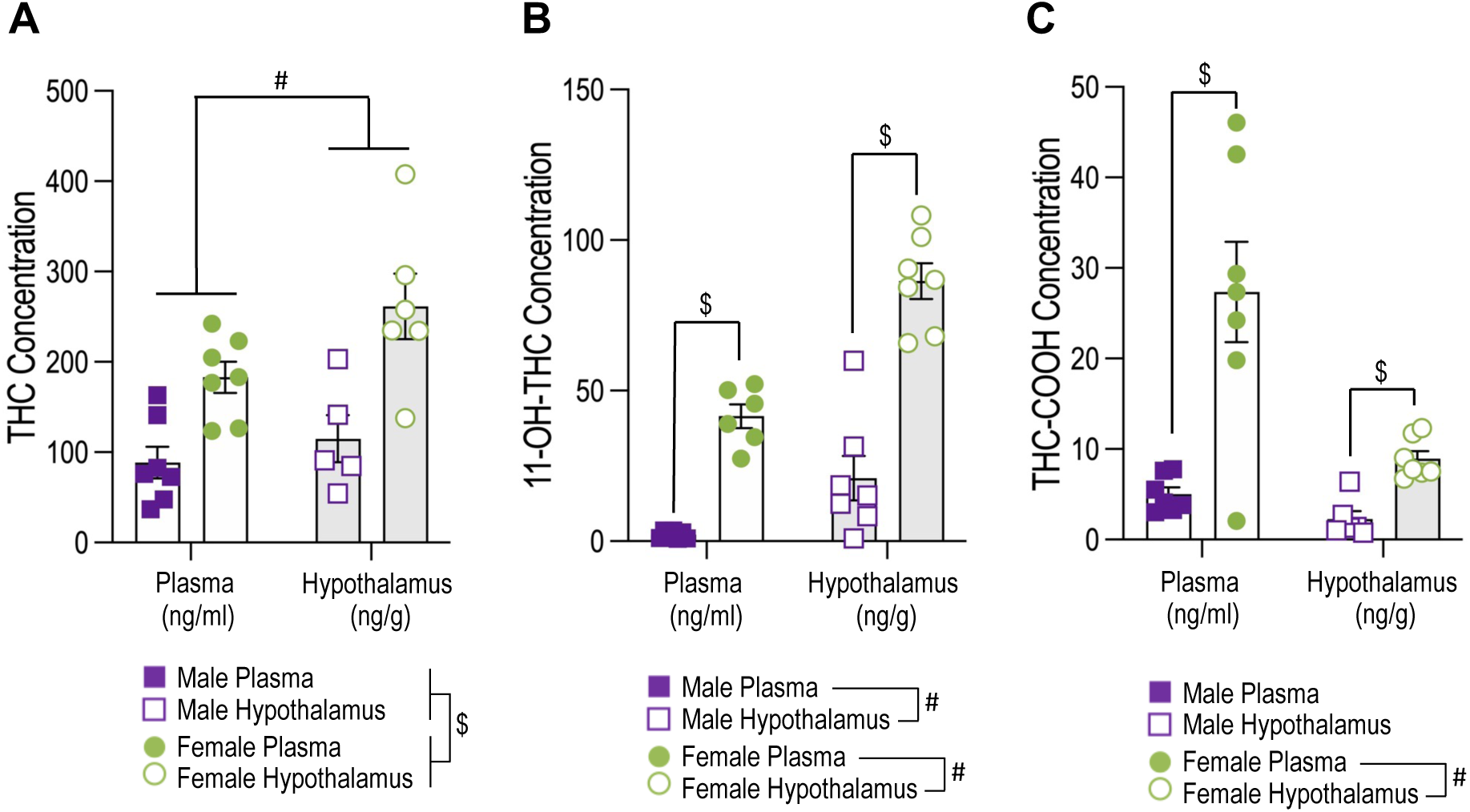
Females have higher THC and metabolite concentrations following THC vapour exposure. **(A)** Mean plasma and hypothalamus concentration of THC (RM 2-way ANOVA, effect of sex: F(1,9)=15.19, p=0.004 ($ females > males); effect of sample: F(1,9)=10.62, p=0.01 (# hypothalamus > plasma)). **(B)** Mean plasma and hypothalamus concentration of 11-OH-THC (RM 2-way ANOVA, interaction effect of sex and sample: F(1,11)=8.58, p=0.014 (post-hoc: $ female > male, # hypothalamus > plasma, p<0.05)). **(C)** Mean plasma and hypothalamus concentration of THC-COOH (RM 2-way ANOVA, interaction effect of sex and sample: F(1,11)=7.79, p=0.018 (post-hoc: $ female > male, # plasma > hypothalamus, p<0.05)). Data shown as mean ± SEM. N=5-7 in each group (two male and one female hypothalamus THC data points, one female plasma 11-OH-THC data point, and one male hypothalamus THC-COOH data point were excluded due to issues with sample processing). # effect of sample, $ effect of sex.

### In the satiated state, THC vapour acutely drives chow intake by increasing feeding bout frequency and duration

To establish if THC vapour can promote food intake in the satiated state, rats were given access to a palatable preload to induce satiety before THC or vehicle vapour exposure, and post-vapour chow intake measured. Prior to vapour exposure, preload consumption significantly supressed chow consumption (effect of preload: F(1,52)=37.68, p<0.001) [Supplementary Figure 1], demonstrating the satiating effect of the preload.

In satiated rats, THC significantly increased chow intake in the first 60-minutes after vapour exposure (interaction of time and treatment: F(2.72,65.26)=30.67, p<0.001) [Figure 3A-C], and this effect is consistent between sexes and increases over exposure days (interaction of time and treatment: F(2,48)=5.96, p=0.005) [Figure 3C]. Vehicle groups continued to demonstrate the satiating effects of the preload during this time [Figure 3A-C]. THC acutely altered feeding patterns in the satiated state [Figure 3D **&** Table 3] by significantly reducing the latency to eat (effect of treatment: F(1,24)=149.2, p<0.0001), increasing feeding bout number (effect of treatment: F(1,24)=44.86, p<0.0001) and duration (effect of treatment: F(1,24)=56.02, p<0.0001) to increase time spent eating (effect of treatment: F(1,24)=56.43, p<0.0001) [Table 3]. THC also increased ingestion rate (effect of treatment: F(1,17)=5.268, p=0.04) [Table 3] and decreased satiety ratio (effect of treatment: F(1,17)=18.63, p=0.0005) [Table 3]. Furthermore, there were no sex differences observed in the effects of THC on feeding parameters [Table 3].

**Figure 3.**
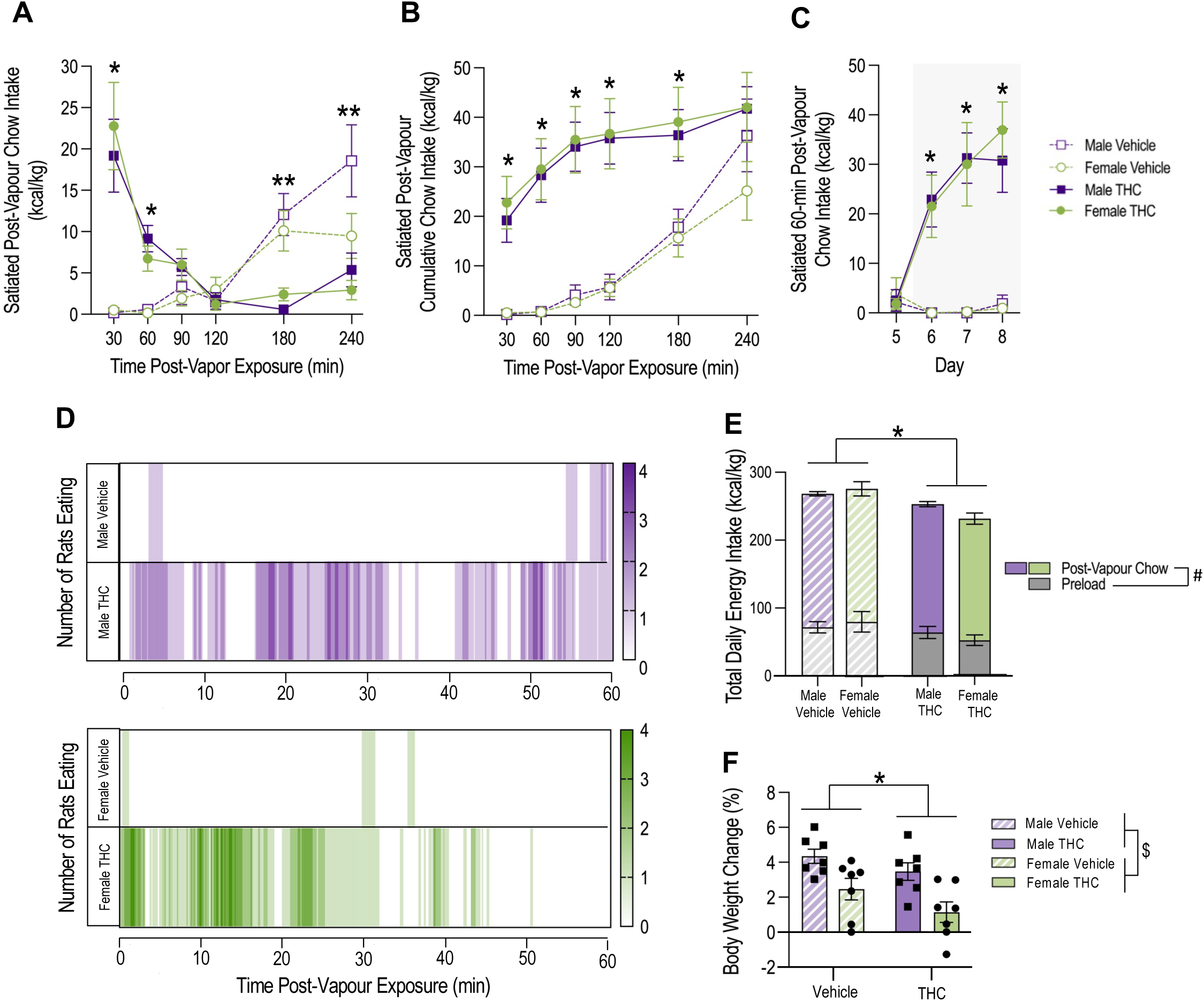
In the satiated state, THC acutely increases chow intake in the first 60-minutes after vapour exposure. **(A)** Day 6-8 mean post-vapour chow intake over time for satiated THC and vehicle groups (RM 3-way ANOVA, interaction effect of time and treatment: F(2.72,65.26)=30.67, p<0.001 (post-hoc: * THC > vehicle at t=30 & 60-minutes, ** vehicle > THC at t=180 & 240-minutes, p<0.01)). **(B)** Day 6-8 mean post-vapour cumulative chow intake for satiated THC and vehicle groups (RM 3-way ANOVA, interaction effect of time and treatment: F(1.9,38.36)=13.30, p<0.001 (post-hoc: * THC > vehicle at t=30, 60, 90, 120 & 180-minutes, p<0.001)). **(C)** Mean 60-minute chow intake for day 5 (baseline, no vapour exposure) and days 6-8 (post-vapour exposure; highlighted in grey) for satiated THC and vehicle groups (RM 3-way ANOVA, interaction effect of time and treatment: F(2.15,51.61)=26.74, p<0.001 (post-hoc: * THC > vehicle on days 6-8, p<0.001)). **(D)** Representative heat maps showing the number and pattern of satiated rats eating chow within the first 60-minutes after vapour exposure for male, female, THC and vehicle groups. **(E)** Day 6-8 mean 24-hour energy intake for THC and vehicle groups (RM 3-way ANOVA, effect of treatment: F(1,24)=273.05, p<0.001 (post-hoc: * THC < vehicle (p<0.01)), effect of energy source: F(1,24)=10.04, p=0.004 (post-hoc: # preload < chow (p<0.001)). **(F)** Body weight change (%) across vapour exposure days (day 6-8) (2-way ANOVA; effect of treatment: F(1,24)=4.3, p=0.049 (* vehicle > THC); effect of sex: F(1,24)=15.53, p<0.001 ($ male > female)). Data normalized to body weight and shown as mean ± SEM. N=7 in each group. * effect of treatment, # effect of food type.

**Table 3.** In satiated conditions, THC acutely increases chow intake by decreasing latency to eat and increasing bout frequency and duration to increase overall time spent eating. Day 4-6 mean feeding behaviour parameters measured within the first 60-minutes after vapour exposure for satiated groups. Data shown as mean ± SEM and analysed using a two-way ANOVA. N=7 in each group, except from ingestion rate and satiety ratio (male vehicle n=3, female vehicle n=4, male & female THC n=7) - these parameters can’t be calculated from animals that don’t eat. Feeding bouts: arbitrarily defined periods of eating separated by 30-seconds or more. Ingestion rate: chow calories consumed / time spent eating. Satiety ratio: time spent not eating / chow calories consumed.

| *Measure* | *SATIATED MALE* |  | *SATIATED FEMALE* |  | *Effect* | *F value* | *P value* |
| --- | --- | --- | --- | --- | --- | --- | --- |
|  | *Vehicle* | *THC* | *Vehicle* | *THC* |  |  |  |
| Chow Intake (kcal) | 0.3 ± 0.3 | 12.2 ± 2.5 | 0.1 ± 0.1 | 6.7 ± 1.4 | Treatment:<br>THC > Vehicle | F(1,24)=41.64 | <0.0001 |
| Time Spent Eating (min) | 0.3 ± 0.2 | 8.0 ± 0.7 | 0.2 ± 0.1 | 7.3 ± 1.9 | Treatment:<br>THC > Vehicle | F(1,24)=56.43 | <0.0001 |
| Latency to Eat (min) | 55.6 ± 2.7 | 13.5 ± 3.1 | 53.1 ± 3.5 | 10.7 ± 4.3 | Treatment:<br>Vehicle > THC | F(1,24)=149.2 | <0.0001 |
| Number of Feeding Bouts | 0.2 ± 0.1 | 4.8 ± 0.4 | 0.2 ± 0.1 | 5.6 ± 1.4 | Treatment:<br>THC > Vehicle | F(1,24)=44.86 | <0.0001 |
| Feeding Bout Duration (min) | 0.2 ± 0.1 | 1.9 ± 0.3 | 0.2 ± 0.1 | 1.5 ± 0.3 | Treatment:<br>THC > Vehicle | F(1,24)=56.02 | <0.0001 |
| First Feeding Bout Duration (min) | 0.1 ± 0.1 | 1.3 ± 0.4 | 0.1 ± 0.1 | 1.1 ± 0.3 | Treatment:<br>THC > Vehicle | F(1,24)=19.41 | 0.0002 |
| Interbout Interval (min) | 54.1 ± 3.0 | 12.5 ± 2.5 | 55.1 ± 2.3 | 14.3 ± 3.2 | Treatment:<br>Vehicle > THC | F(1,24)=218.7 | <0.0001 |
| Ingestion Rate (kcal/min) | 0.7 ± 0.4 | 1.9 ± 0.5 | 0.4 ± 0.1 | 1.1 ± 0.1 | Treatment:<br>THC > Vehicle | F(1,17)=5.268 | 0.0347 |
| Satiety Ratio (min/kcal) | 323.4 ± 171.0 | 6.6 ± 1.5 | 268.5 ± 118.6 | 12.4 ± 2.7 | Treatment:<br>Vehicle > THC | F(1,17)=18.63 | 0.0005 |

Like the previous experiment, where rats were exposed to THC in standard conditions, satiated animals showing acute THC-driven feeding reduce subsequent food intake so that there is no significant difference in the total amount of chow eaten 240-minutes following vapour exposure between vehicle and THC groups (no effect of treatment: F(1,24)=3.13, p=0.09) [Figure 3B]. However, total daily energy intake was influenced by THC, where both male and female THC groups consumed less daily energy compared to vehicle groups (effect of treatment: F(1,24)=9.18, p=0.006) [Figure 3E], and this is reflected in the THC groups showing less body weight change across vapour exposure days (effect of treatment: F(1,24)=4.3, p=0.049) [Figure 3F].

### THC vapour acutely increases feeding regardless of food macronutrient content by increasing feeding bout frequency

To determine if the type of food available influences the effects of THC on acute feeding patterns, rats were exposed to THC or vehicle vapour, given access to either a high-carbohydrate or a high-fat food [Table 1] and intake measured.

Regardless of macronutrient content, THC significantly increased food intake in the first 60-minutes following vapour exposure (*high-carbohydrate access:* effect of treatment: F(1,24)=20.41, p<0.001; *high-fat access:* effect of treatment: F(1,24)=18.69, p<0.001) [Figure 4].

**Figure 4.**
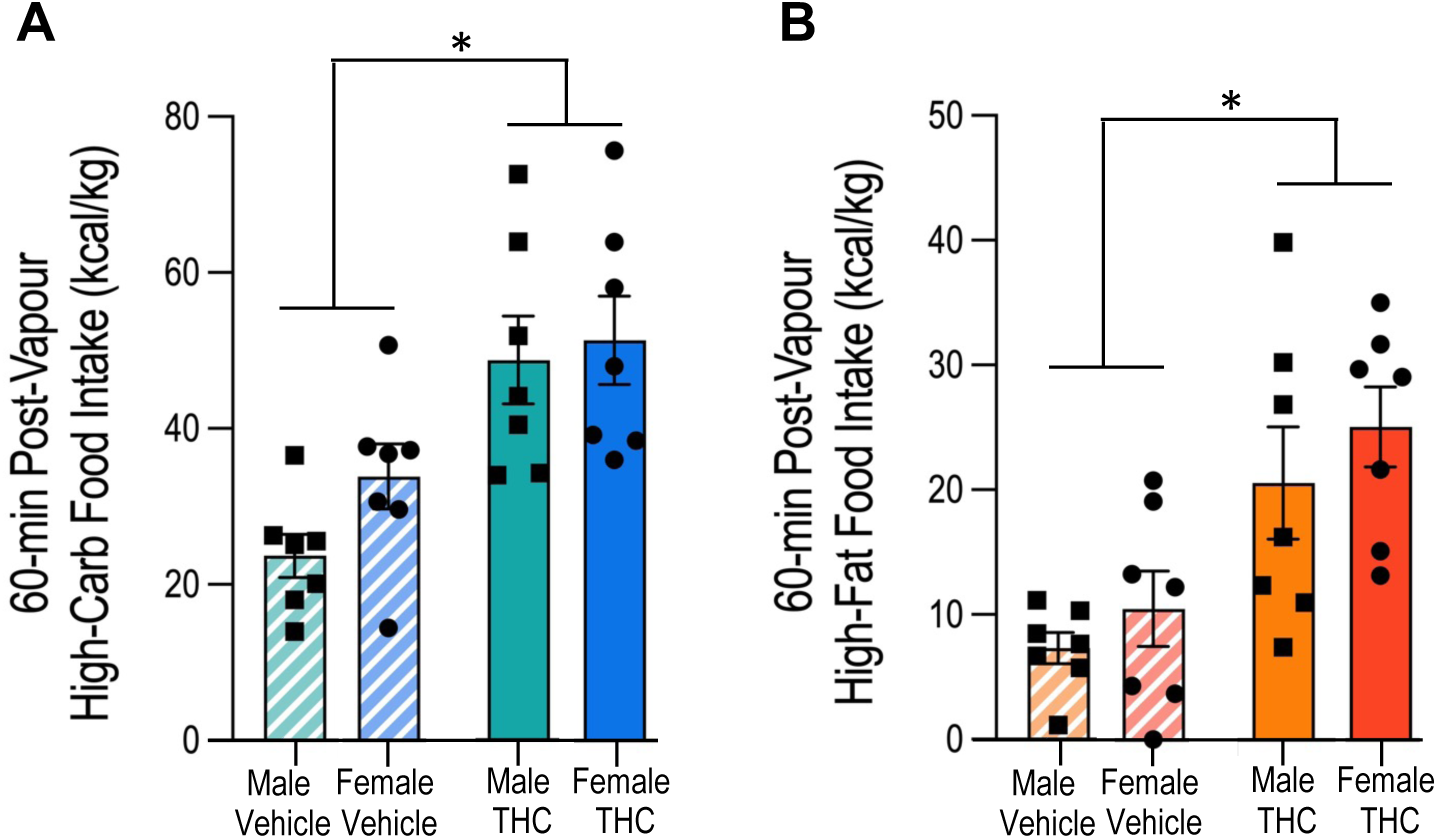
Regardless of food macronutrient content, THC increases intake in the first 60-minutes after vapour exposure. **(A)** Day 6-8 mean 60-minute post-vapour intake of high-carbohydrate food for THC and vehicle groups (2-way ANOVA; effect of treatment: F(1,24)=20.41, p<0.001 (* THC > Vehicle)). **(B)** Day 6-8 mean 60-minute post-vapour intake of high-fat food for THC and vehicle groups (RM 2-way ANOVA; effect of treatment: F(1,24)=18.69, p<0.001 (* THC > Vehicle)). All data normalized to body weight and shown as mean ± SEM. N=7 in each group. * effect of treatment.

Within this time, THC altered feeding patterns of both foods [Supplementary Figure 2D and 3D & Supplementary Table 1] through decreasing latency to eat (*high-carbohydrate access:* effect of treatment: F(1,24)=11.37, p=0.003; *high-fat access*: effect of treatment: F(1,24)=7.39, p=0.012) and increasing bout number (*high-carbohydrate access:* effect of treatment F(1,24)=4.37, p=0.047; *high-fat access*: effect of treatment: F(1,24)=26.75, p<0.001), to increase time spent eating (*high-carbohydrate access:* effect of treatment F(1,24)=12.85, p=0.002; *high-fat access*: effect of treatment: F(1,24)=16.48, p<0.001) [Supplementary Table 1]. With high-carbohydrate food, THC also decreased bout duration (effect of treatment: F(1,24)=21.31, p=0.0001) and satiety ratio (effect of treatment: F(1,24)=15.68, p=0.0006) [Supplementary Table 1]. With high-fat food, THC had no effect on bout duration (no effect of treatment: F(1,24)=0.05, p=0.82) or satiety ratio (no effect of treatment: F(1,21)=2.42, p=0.14), and irrespective of food, THC did not affect ingestion rate (*high-carbohydrate access:* no effect of treatment: F(1,24)=1.97, p=0.17; *high-fat access*: no effect of treatment: F(1,21)=2.77, p=0.11) [Supplementary Table 1]. Furthermore, a sex difference was detected, where THC increased the number high-carbohydrate food feeding bouts to a greater extent in males compared to females (interaction between treatment and sex: F(1,24)=4.37, p=0.047) [Supplementary Table 1]. Irrespective of treatment, males had a higher high-carbohydrate food interbout interval (effect of sex: F(1,24)=4.97, p=0.036) and ingestion rate (effect of sex: F(1,24)=9.47, p=0.005) compared to females [Supplementary Table 1].

With macronutrient-rich food, the timeframe and day-to-day consistency of THC effects on feeding was like that seen in chow access experiments [Supplementary Figure 2A-C & 3A-C]. Also in line with chow experiments, THC did not influence total energy intake in the 240-minute period following vapour exposure (*high-carbohydrate access:* no effect of treatment: F(1,24)=0.01921, p=0.9; *high-fat access:* no effect of treatment: F(1,24)=0.006, p=0.9) [Supplementary Figure 2B & 3B], but THC reduced total daily energy intake only in rats that had post-vapour access to high-carbohydrate food (*high-carbohydrate access*: effect of treatment: F(1,24)=11.01, p=0.003; *high-fat access*: no effect of treatment: F(1,24)=0.94, p=0.34) [Supplementary Figure 2E & 3E]. However, THC had no significant effect on body weight change with either high-carbohydrate or high-fat food access post-vapour (*high-carbohydrate access*: no effect of treatment: F(1,24)=3.77, p=0.064; *high-fat access*: no effect of treatment: F(1,24)=0.068, p=0.8) [Supplementary Figure 2F & 3F]. Irrespective of treatment, a sex difference was detected where females eat overall more high-carbohydrate or high-fat food than males but eat less chow (*high-carbohydrate access*: interaction between food and sex: F(1,24)=7.90, p=0.01; *high-fat access*: interaction between food and sex: F(1,24)=28.04, p<0.001) [Supplementary Figure 2E & 3E], and females gain less body weight over vapour exposure days compared to males (*high-carbohydrate access*: effect of sex: F(1,24)=24.30, p<0.001; *high-fat access*: effect of sex: F(1,24)=27.42, p<0.001) [Supplementary Figure 2F & 3F].

### THC vapour acutely and selectively increases high-fat food intake, abolishing pre-existing high-carbohydrate food preference

To investigate if THC vapour alters macronutrient-specific food preferences, rats were exposed to THC or vehicle vapour, given access to both a high-carbohydrate and a high-fat food [Table 1] and intake of each food simultaneously measured.

With vehicle treatment (and at baseline), rats had a significant preference for the high-carbohydrate food (vehicle high-carbohydrate preference > vehicle high-fat preference: F(1,12)=17.61, p=0.001) [Figure 5A&B]. However, after THC vapour exposure both male and female rats selectively increased the amount of high-fat food eaten within the first 60-minutes after vapour exposure (THC high-fat intake > vehicle high-fat intake: F(1,24)=8.25, p=0.008) [Figure 5A]. Thereby, THC significantly increased high-fat food preference (THC high-fat preference > vehicle high-fat preference: F(1,24)=10.21, p=0.004) and consequently animals lost their high-carbohydrate food preference and no longer significantly preferred one food over the other (no difference between high-fat and high-carbohydrate food preferences: F(1,12)=1.07, p=0.32) [Figure 5B]. Irrespective of treatment, females generally had a higher preference for the high-fat food compared to males (sex difference: F(1,24)=4.56, p=0.043) [Figure 5B].

**Figure 5.**
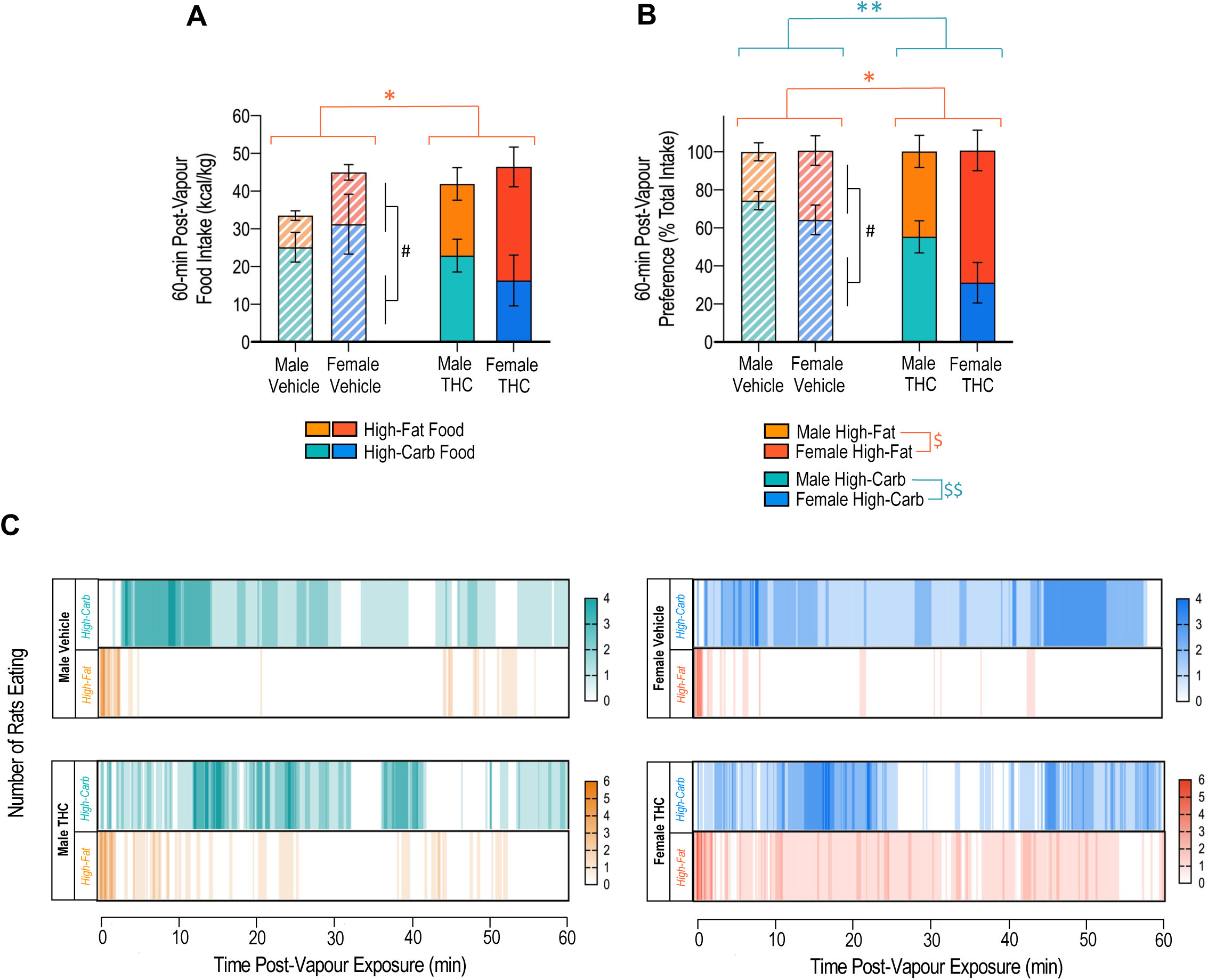
When given a food choice, THC acutely and selectively increases high-fat food intake to abolish pre-existing high-carbohydrate food preference. **(A)** Day 6-8 mean 60-minute post-vapour intake of high-carbohydrate and high-fat food for THC and vehicle groups (RM 3-way ANOVA; interaction effect of vapour treatment and food type: F(1,24)=8.25, p=0.008 (post-hoc: * THC high-fat intake > vehicle high-fat intake, # vehicle high-carbohydrate intake > vehicle high-fat intake, p=0.01)). **(B)** Day 6-8 mean 60-minute post-vapour high-carbohydrate and high-fat food preference (% total intake) for THC and vehicle groups (RM 3-way ANOVA; interaction effect of vapour treatment and food type: F(1,24)=10.21, p=0.004 (post-hoc: * THC high-fat food preference > vehicle high-fat food preference, ** THC high-carbohydrate food preference < vehicle high-carbohydrate food preference, # vehicle high-carbohydrate food preference > vehicle high-fat food preference, p<0.01), interaction effect of food type and sex: F(1,24)=4.56, p=0.043 (post-hoc: $ female high-fat food preference > male high-fat food preference, $$ female high-carbohydrate food preference < male high-carbohydrate food preference, p<0.05)). **(C)** Representative heat maps showing the number and pattern of rats eating high-carbohydrate and high-fat food across the first 60-minutes after vapour exposure. Food intake data normalized to body weight and shown as mean ± SEM. N=7 in each group. * effect of treatment, $ effect of sex, # effect of food type.

Interestingly, THC did not significantly alter the latency to eat either food (no effect of treatment: F(1,24)=0.06, p=0.82) [Table 4] or feeding latency in general (no effect of treatment: F(1,24)=0.02, p=0.3). However, THC did alter feeding patterns [Figure 5C **&** Table 4] in the first 60-minutes after vapour exposure by significantly increasing feeding bout number irrespective of food (effect of treatment: F(1,24)=14.91, p<0.001) [Table 4] and decreasing high-carbohydrate food feeding bout duration (interaction of food and treatment: F(1,24)=13.72, p=0.001) [Table 4]. Rats generally spent more time eating the high-carbohydrate food (effect of food: F(1,24)=10.68, p=0.003) [Table 4], but a trend was noted where THC increased time spent eating the high-fat food, however this did not reach significance [Table 4]. Furthermore, THC decreased ingestion rate for high-carbohydrate food, particularly in females (interaction of food, treatment, and sex: F(1,21)=4.62, p=0.043) [Table 4], and THC had no effect on satiety ratio (no effect of treatment: F(1,21)=2.10, p=0.16) [Table 4].

**Table 4.** When given a food choice, THC increases feeding bout frequency irrespective of food type, but decreases high-carbohydrate feeding bout duration, thereby THC increases time spent eating high-fat food but does not alter time spent eating high-carbohydrate food. Day 6-8 mean feeding behaviour parameters measured within the first 60-minutes after vapour exposure. Data shown as mean ± SEM and analysed using a three-way ANOVA (for parameters with no effect, only the F and P values for treatment comparison are shown). N=7 in each group except from ingestion rate and satiety ratio (male THC n=6, female vehicle n=5, male vehicle & female THC n=7) - these parameters can’t be calculated and compared from animals that don’t eat both foods. Feeding bouts: arbitrarily defined periods of eating separated by 30-seconds or more. Ingestion rate: chow calories consumed / time spent eating. Satiety ratio: time spent not eating / chow calories consumed. High-carb: high-carbohydrate food. High-fat: high-fat food.

| Measure | MALE |  |  |  | FEMALE |  |  |  | Effect | F Value | P Value |
| --- | --- | --- | --- | --- | --- | --- | --- | --- | --- | --- | --- |
|  | High-Carbohydrate |  | High-Fat |  | High-Carbohydrate |  | High-Fat |  |  |  |  |
|  | Vehicle | THC | Vehicle | THC | Vehicle | THC | Vehicle | THC |  |  |  |
| Intake (kcal) | 11.6 ± 1.8 | 10.5 ± 2.7 | 3.7 ± 0.7 | 8.7 ± 2.9 | 10.2 ± 2.5 | 4.1 ± 1.6 | 3.7 ± 1.0 | 8.5 ± 2.7 | Food x Treatment:<br>THC > Vehicle<br>Vehicle High-Carb > Vehicle High-Fat | F(1,24)=6.64 | 0.017 |
| Time Spent Eating (min) | 11.0 ± 2.5 | 10.3 ± 2.7 | 1.5 ± 0.2 | 3.0 ± 1.0 | 13.8 ± 2.7 | 10.4 ± 3.1 | 1.0 ± 0.2 | 9.7 ± 6.2 | Food:<br>High-Carb > High-Fat | F(1,24)=10.68 | 0.003 |
| Latency to Eat (min) | 14.8 ± 7.7 | 9.6 ± 5.6 | 0.9 ± 0.3 | 8.1 ± 7.0 | 2.4 ± 1.1 | 4.9 ± 2.2 | 7.1 ± 6.0 | 0.5 ± 0.1 | No effect of treatment/sex/food | F(1,24)=0.06 | 0.816 |
| Number of Feeding Bouts | 2.6 ± 0.6 | 8.1 ± 2.6 | 3.3 ± 0.5 | 4.6 ± 1.3 | 3.4 ± 0.4 | 8.7 ± 1.4 | 2.7 ± 0.5 | 8.1 ± 2.3 | Treatment:<br>THC > Vehicle | F(1,24)=14.91 | <0.001 |
| Feeding Bout Duration (min) | 4.9 ± 0.8 | 1.8 ± 0.7 | 0.5 ± 0.1 | 0.6 ± 0.2 | 4.4 ± 1.0 | 1.7 ± 0.6 | 0.3 ± 0.1 | 1.1 ± 0.6 | Food x Treatment:<br>Vehicle > THC<br>Vehicle High-Carb > Vehicle High-Fat | F(1,24)=13.72 | 0.001 |
| First Feeding Bout Duration (min) | 4.4 ± 1.4 | 1.0 ± 0.7 | 0.7 ± 0.2 | 0.8 ± 0.2 | 3.2 ± 2.0 | 0.4 ± 0.2 | 0.5 ± 0.1 | 0.6 ± 0.2 | Food x Treatment:<br>Vehicle > THC<br>Vehicle High-Carb > Vehicle High-Fat | F(1,24)=6.16 | 0.020 |
| Interbout Interval (min) | 20.1 ± 6.1 | 10.9 ± 3.8 | 14.7 ± 1.5 | 15.0 ± 4.0 | 10.7 ± 1.3 | 5.9 ± 1.0 | 18.3 ± 3.1 | 8.9 ± 2.5 | Treatment:<br>Vehicle > THC | F(1,24)=6.27 | 0.019 |
| Ingestion Rate (kcal/min) | 1.3 ± 0.3 | 0.9 ± 0.2 | 2.3 ± 0.3 | 3.6 ± 0.9 | 0.6 ± 0.1 | 0.3 ± 0.1 | 7.1 ± 3.1 | 2.5 ± 0.8 | Food x Treatment x Sex:<br>Female Vehicle High-Carb > Female THC High-Carb<br>Female Vehicle High-Fat > Male Vehicle High-Fat<br>Male THC High-Carb > Female THC High-Carb<br>Female Vehicle High-Fat > Female Vehicle High-Carb | F(1,21)=4.62 | 0.043 |
| Satiety Ratio (min/kcal) | 5.2 ± 1.2 | 4.8 ± 0.9 | 30.2 ± 15.4 | 27.1 ± 16.5 | 4.2 ± 0.7 | 52.0 ± 21.6 | 16.8 ± 3.2 | 14.8 ± 6.5 | No effect of treatment/sex/food | F(1,21)=2.10 | 0.163 |

Furthermore, the timeframe and day-to-day consistency of THC effects was like previous experiments [Supplementary Figure 4A-C] and THC decreased total daily energy intake (effect of treatment: F(1,24)=10.43, p=0.004) [Supplementary Figure 4D], but this was not reflected in changes in body weight (no effect of treatment: F(1,24)=3.41, p=0.08) [Supplementary Figure 4E].

### In the satiated state, THC vapour acutely increases intake of both high-carbohydrate and high-fat foods, abolishing pre-existing high-fat food preference

To assess if THC vapour can alter macronutrient-specific food preferences in the satiated state, rats were satiated through the consumption of a palatable preload before being exposed to either THC or vehicle vapour, then given access to both a high-carbohydrate and a high-fat food [Table 1] and intake of each food measured simultaneously.

At baseline, preload consumption significantly reduced high-carbohydrate and high-fat food consumption (preload < no preload: F(1,50)=70.65, p<0.001) [Supplementary Figure 5], demonstrating the satiating effect of the preload on high-carbohydrate and high-fat food intake.

With vehicle treatment (and at baseline), satiated rats showed a significant preference for the high-fat food (vehicle high-fat food preference > vehicle high-carbohydrate food preference: F(1,8)=16.83, p=0.003) [Figure 6A&B]. However, THC vapour exposure significantly increased intake of both the high-carbohydrate and high-fat foods within the first 60-minutes after vapour exposure (THC intake > vehicle intake: F(1,22)=19.86, p<0.001) [Figure 6A], where females generally ate more than males (female intake > male intake: F(1,22)=6.83, p<0.016) [Figure 6A]. This resulted in THC exposed animals losing their high-fat food preference, no longer significantly preferring one food over the other (no difference between high-fat and high-carbohydrate food preferences: F(1,12)=0.91, p=0.36) [Figure 6B].

**Figure 6.**
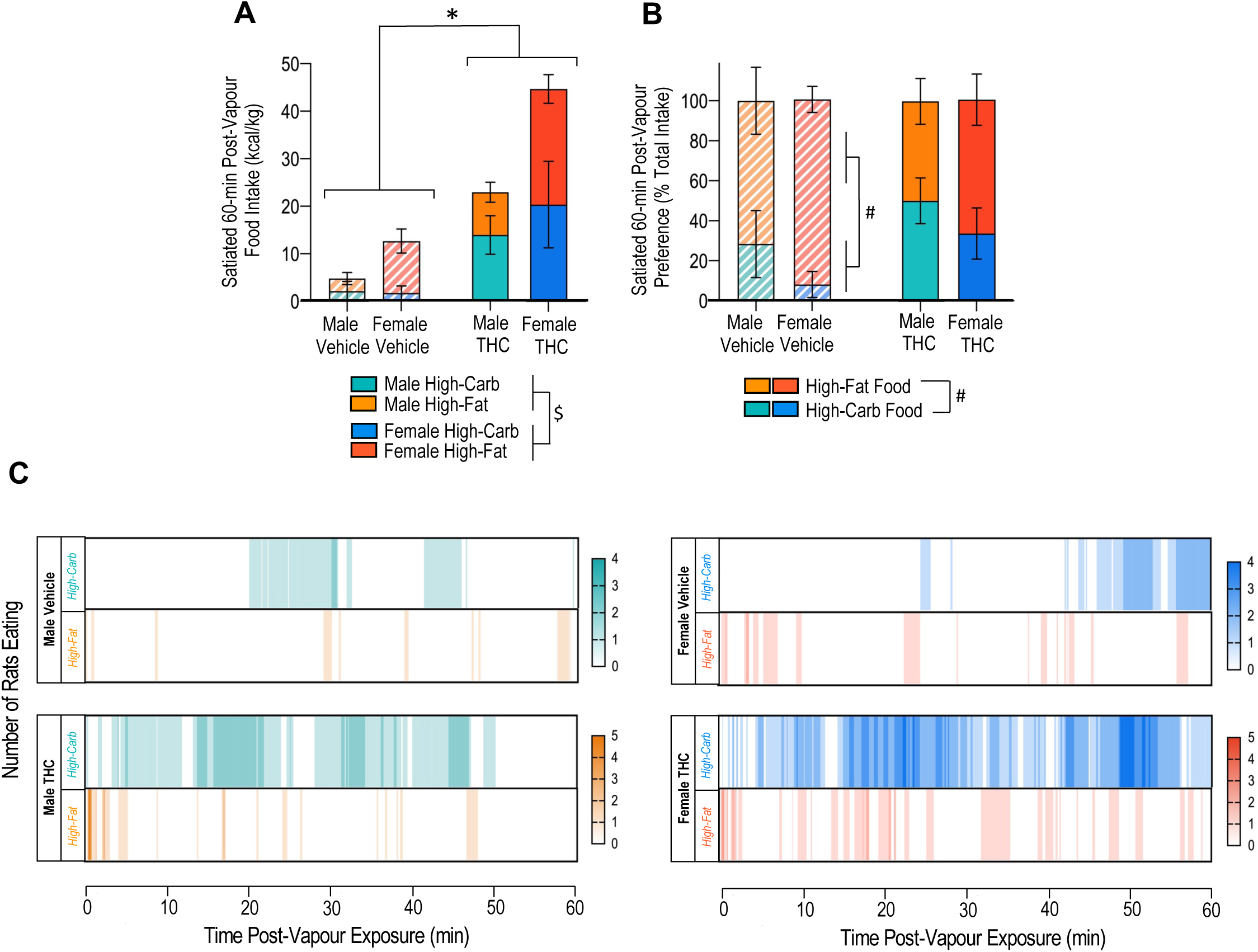
In satiated conditions, when given a food choice, THC acutely increases intake of both high-carbohydrate and high-fat food, but overall dampens pre-existing high-fat food preference. **(A)** Day 6-8 mean 60-minute post-vapour intake of high-carbohydrate and high-fat food for THC (n=7 in each group) and vehicle (n=6 in each group) groups (RM 3-way ANOVA; effect of treatment: F(1,22)=19.86, p<0.001 (* THC > vehicle); effect of sex: F(1,22)=6.83, p<0.016 ($ female > male)). **(B)** Day 6-8 mean 60-minute post-vapour high-carbohydrate and high-fat food preference (% total intake) for THC (n=7 in each group) and vehicle (male vehicle: n=4, female vehicle: n=6) groups (RM 2-way ANOVA; *vehicle:* # effect of food type: F(1,8)=16.83, p=0.003; THC: no effect of food: F(1,12)=0.91, p=0.35). **(D)** Representative heat maps showing the number and pattern of rats eating high-carbohydrate and high-fat food 60-minutes post-vapour exposure. Food intake data normalized to body weight and shown as mean ± SEM. One rat in each the male and female vehicle group were excluded based on meeting outlier criteria (>2.5 SD away from the mean). Data from two other male vehicle rats could not be included the preference analysis as they did not eat any food in the first 60-minutes after vapour exposure, therefore, preference could not be assessed. * effect of treatment, $ effect of sex, # effect of food type.

THC altered feeding patterns [Figure 6C **&** Table 5] through decreasing latency to eat the high-carbohydrate food (interaction of food and treatment: F(1,22)=7.31, p=0.013) [Table 5], increasing the number of feeding bouts irrespective of food (effect of treatment: F(1,22)=18.68, p<0.001) [Table 5], and increasing high-carbohydrate food bout duration (interaction of food and treatment: F(1,22)=6.90, p=0.02) [Table 5], to increase time spent eating the high-carbohydrate food (interaction of food and treatment: F(1,22)=4.35, p=0.049) [Table 5]. Other behavioural parameters remained unchanged after THC exposure, including ingestion rate (no effect of treatment: F(1,9)=0.16, p=0.7) and satiety ratio (no effect of treatment: F(1,9)=0.41, p=0.54) [Table 5]. Irrespective of treatment, a sex difference was detected where females generally showed a greater number of feeding bouts compared to males (effect of sex: F(1,22)=4.51, p=0.045).

**Table 5.** In satiated conditions, when given a food choice, THC promotes food intake by decreasing latency to eat high-carbohydrate food, increasing feeding bout frequency and duration to increase overall time spent eating the high-carbohydrate food. Day 6-8 mean feeding behaviour parameters measured within the first 60-minutes after vapour exposure. Data shown as mean ± SEM and analysed using a two-way ANOVA. N=7 in each group, except from ingestion rate and satiety ratio (male vehicle n=3, female vehicle n=4, male & female THC n=7) - these parameters can’t be calculated from animals that don’t eat. Feeding bouts: arbitrarily defined periods of eating separated by 30-seconds or more. Ingestion rate: chow calories consumed / time spent eating. Satiety ratio: time spent not eating / chow calories consumed. High-carb: high-carbohydrate food. High-fat: high-fat food.

| Measure | SATIATED MALE |  |  |  | SATIATED FEMALE |  |  |  | Effect | F Value | P Value |
| --- | --- | --- | --- | --- | --- | --- | --- | --- | --- | --- | --- |
|  | High-Carbohydrate |  | High-Fat |  | High-Carbohydrate |  | High-Fat |  |  |  |  |
|  | Vehicle | THC | Vehicle | THC | Vehicle | THC | Vehicle | THC |  |  |  |
| Intake (kcal) | 2.3 ± 1.5 | 6.7 ± 3.4 | 0.7 ± 0.4 | 2.5 ± 0.8 | 3.5 ± 2.5 | 8.0 ± 3.3 | 2.4 ± 0.8 | 5.3 ± 1.1 | Treatment:<br>THC > Vehicle | F(1,22)=8.96 | 0.007 |
| Time Spent Eating (min) | 2.3 ± 1.6 | 7.5 ± 3.5 | 0.6 ± 0.3 | 1.1 ± 0.3 | 3.2 ± 2.1 | 13.5 ± 4.0 | 1.2 ± 0.3 | 2.7 ± 0.6 | Food x Treatment:<br>THC High-Carb > Vehicle High-Carb<br>Vehicle High-Carb > Vehicle High-Fat | F(1,22)=4.35 | 0.049 |
| Latency to Eat (min) | 49.9 ± 6.5 | 20.4 ± 9.4 | 28.3 ± 9.8 | 9.3 ± 8.5 | 47.9 ± 6.0 | 11.0 ± 4.1 | 19.2 ± 8.3 | 11.3 ± 7.7 | Food x Treatment:<br>THC High-Carb < Vehicle High-Carb<br>THC High-Carb > THC High-Fat | F(1,22)=7.31 | 0.013 |
| Number of Feeding Bouts | 0.9 ± 0.5 | 2.6 ± 0.7 | 1.3 ± 0.5 | 3.0 ± 0.7 | 1.1 ± 0.6 | 6.7 ± 1.9 | 2.3 ± 0.5 | 4.3 ± 0.7 | Treatment:<br>THC > Vehicle | F(1,22)=18.68 | <0.001 |
|  |  |  |  |  |  |  |  |  | Sex:<br>Female > Male | F(1,22)=4.51 | 0.045 |
| Feeding Bout Duration (min) | 1.0 ± 0.8 | 2.2 ± 0.9 | 0.3 ± 0.1 | 0.4 ± 0.1 | 0.9 ± 0.6 | 1.9 ± 0.5 | 0.6 ± 0.2 | 0.7 ± 0.2 | Food x Treatment:<br>THC High-Carb > Vehicle High-Carb<br>THC High-Carb > THC High-Fat | F(1,22)=6.90 | 0.015 |
| First Feeding Bout Duration (min) | 1.5 ± 1.4 | 1.5 ± 1.1 | 0.3 ± 0.1 | 0.3 ± 0.1 | 0.2 ± 0.1 | 0.2 ± 0.1 | 0.4 ± 0.2 | 0.4 ± 0.1 | No effect of treatment/sex/food | F(1,22)=0.98 | 0.334 |
| Interbout Interval (min) | 47.1 ± 8.3 | 22.5 ± 7.2 | 34.4 ± 7.1 | 20.5 ± 6.7 | 42.6 ± 8.4 | 10.4 ± 3.7 | 24.1 ± 5.8 | 12.0 ± 1.6 | Food x Treatment:<br>Vehicle High-Carb > THC High-Carb<br>Vehicle High-Fat > THC High-Fat<br>Vehicle High-Carb > Vehicle High-Fat | F(1,22)=5.53 | 0.028 |
| Ingestion Rate (kcal/min) | 0.9 ± 0.4 | 0.6 ± 0.3 | 2.4 ± 1.8 | 2.2 ± 0.7 | 0.7 ± 0.4 | 0.4 ± 0.1 | 2.1 ± 0.6 | 2.4 ± 0.5 | No effect of treatment/sex/food | F(1,9)=0.16 | 0.699 |
| Satiety Ratio (min/kcal) | 529.8 ± 523.3 | 168.5 ± 154.1 | 42.9 ± 30.5 | 96.1 ± 46.2 | 5.0 ± 2.3 | 253.2 ± 248.6 | 38.5 ± 17.3 | 12.7 ± 1.7 | No effect of treatment/sex/food | F(1,9)=0.41 | 0.537 |

Moreover, the timeframe and day-to-day consistency of THC effects was similar to previous experiments) [Supplementary Figure 6A-C], and THC had no effect on total daily energy intake (no effect of treatment: F(1,22)=0.56, p=0.46) [Supplementary Figure 6D], or body weight change (no effect of treatment: F(1,22)=0.13, p=0.72) [Supplementary Figure 6E].

## Discussion

The effects of cannabis inhalation on feeding behaviours have not been fully explored, therefore the aim of this study was to use a translational pre-clinical rat model to fully characterise how inhalation of vapour from a THC-dominant cannabis extract influences food intake and choice. To our knowledge this is the first study to determine how THC vapour inhalation alters feeding using microstructural feeding pattern analysis, the first to investigate if sex differences in THC vapour inhalation driven feeding behaviours exists, and the first to directly measure how THC alters macronutrient-specific food preferences using a free choice paradigm.

We found that THC vapour inhalation robustly and acutely increased food intake regardless of sex, the macronutrient content or palatability of food available, or whether animals are satiated or not. Consistently, we showed that THC vapour inhalation reduced latency to eat, and increased feeding bout number to increase food intake. Throughout the study, these acute appetitive THC effects were compensated for through reductions in subsequent food intake, thereby in most experiments, body weight change was not significantly affected. Furthermore, we provide direct evidence that THC can alter macronutrient-specific food preferences, where when given a choice between a high-fat and a high-carbohydrate food, THC vapour increases fat preference in regular conditions, and trends towards increasing carbohydrate preference in satiated conditions, so that animals no longer specifically prefer one food over the other. Finally, no notable sex differences in THC effects were observed in experiments where rats were given a single food following THC vapour exposure, however, when given a food choice, females responded to THC with a stronger preference for the high-fat food compared to males.

Our model consistently showed that THC vapour inhalation increases food intake in the first 60-minutes after exposure. This timeframe is in line with the pharmacokinetic profile of THC vapour inhalation when administered using our protocol, where peak blood THC concentrations are detected immediately after vapour exposure and brain THC concentrations are highest throughout the first 60-minutes following vapour exposure [35]. In addition, the time course of our THC feeding effects are consistent with that of previous THC or cannabis vapour exposure studies [19, 23].

We undertook microstructural feeding pattern analysis to determine how THC vapour inhalation acutely influences behaviour to increase food intake, as this has not been previously undertaken. THC vapour could alter food intake by altering the initiation, termination, frequency, and/or rate of feeding, thereby we measured the latency to eat, feeding bout number and duration, interbout interval, eating rate and satiety ratio. Consistently, THC vapour inhalation decreased the latency to eat, and this was not dependent on sex, food type or satiety. This indicates that THC increases feeding motivation, supported by inhalation studies directly measuring motivation to acquire food [16]. These motivation effects are not specific to route of administration as oral and injected THC have also been shown to decrease feeding latency [26, 44], and increase motivation to acquire food [15, 17] respectively. However, in our standard food choice experiment, latency to eat was unaffected by THC vapour exposure, presumably due to all animals being highly motivated to eat when given access to multiple palatable foods. Yet, in the satiated food choice experiment, we saw a robust effect of THC vapour in reducing latency to eat, as consumption of the preload likely reduces baseline motivation for these palatable foods.

Furthermore, we show that THC vapour exposed rats do not binge eat all food in a single bout. In all experiments, THC vapour increased the number of feeding bouts, but only increased feeding bout duration in satiated conditions. Thereby, rats exposed to THC vapour in standard conditions increase food intake by eating more frequently. Satiated THC vapour exposed animals eat more frequently and spend more time eating within each feeding bout, consistent with satiated oral THC administration [25, 44]. Differences between satiated and standard conditions can most likely be explained through the effects of satiety on baseline feeding patterns.

Additionally, it doesn’t appear that THC vapour consistently alters feeding by changing consumption rate, as in most experiments, ingestion rate remained unchanged. However, THC vapour significantly increased the rate of chow consumption in the satiated state and significantly reduced the rate of high-carbohydrate food consumption in the standard food choice experiment. Thereby, THC may alter ingestion rate in certain situations involving satiety or palatable foods, however, these findings were not consistent between our experiments.

Moreover, in experiments that did not involve high-fat food access, THC vapour significantly decreased satiety ratio compared to vehicle groups demonstrating that THC vapour can reduce the satiating potential of food eaten. Experiments giving high-fat food access showed no effect of THC vapour on satiety ratio, potentially due to high-fat food being more satiating in general compared to other foods.

We also show that rats compensate for this acute THC-induced hyperphagic response by reducing subsequent food intake, so that total energy intake in the 240-minute period after vapour exposure is no different between THC and vehicle groups. Similar compensatory effects were observed in previous THC vapour experiments [19], as well as in response to low doses of oral or injected THC [10, 20]. Interestingly, these compensatory effects were apparent regardless of sex and the type of food given, demonstrating how robust acute homeostatic responses to palatable, energy-dense foods are in rats.

Effects of THC vapour exposure on 24-hour energy intake were variable between experiments, with some showing that THC vapour decreases daily energy intake and others having no significant effect. These between-experiment differences were not associated with the state of the animals or food type. Furthermore, most experiments showed that THC vapour exposure does not significantly change body weight gain, suggesting that these short-term reductions in daily food intake are not significant enough to influence body weight, but with a longer THC exposure, body weight differences may become more apparent.

We found that THC vapour exposure can drive food intake in the satiated state, whether given chow or a palatable food choice. Similar effects have been reported in studies administering oral THC [10, 25, 26] or vaporised cannabis [23]. Thus, THC may override homeostatic satiety mechanisms to stimulate feeding. Furthermore, THC vapour inhalation can increase the motivation to acquire chow [16] and injected THC increases hedonic orofacial responses to sucrose [21, 22] or aversive bitter solutions [22]. Suggesting that THC might increase the palatability and/or reward value of foods to drive intake, even in situations where the food is devalued i.e., when aversive or during satiety.

To our knowledge this is the first study to directly measure how THC alters macronutrient-specific food preferences in rats. Most previous literature implies that THC increases preference for a high-carbohydrate food, where regular cannabis use is associated with higher consumption of carbohydrates [24] and sweet snack foods [14]. Furthermore, in rodents, injection of a cannabinoid 1 receptor (CB1R) agonist increases carbohydrate intake when given a choice between carbohydrate, protein, and fat [45], and injection of a CB1R antagonist selectively reduces high-sucrose chow intake when given a choice between high-sucrose and regular chow [46, 47]. However, in this study we show that THC vapour inhalation in regular conditions has no effect on high-carbohydrate food intake, but selectively increases high-fat food intake. Thereby, in contrast to the evidence outlined above, our model shows that THC increases high-fat food preference. A small body of evidence aligns with this finding, where following injected THC, rats with access to a high-fat diet eat more than rats with access to a sweetened high-fat diet [20], and injection of a CB1R antagonist dampens high-fat diet associated place preference in mice [48]. Our microstructural feeding pattern analysis shows that THC increases the number of feeding bouts to increase time spent eating the high-fat food. THC did not alter time spent eating the high-carbohydrate food as increases in the number of feeding bouts were offset by decreased feeding bout duration. Furthermore, in our choice paradigm, the high-fat food contains more energy than the high-carbohydrate food, therefore, THC vapour may also be inducing preference for energy density. Yet, in satiated conditions, we showed that THC vapour increased intake of both high-carbohydrate and high-fat foods, but trended towards increasing high-carbohydrate food preference, supporting previous literature [45–47]. In this experiment, THC specifically reduced the latency to eat high-carbohydrate food, implying that rats were more motivated to eat this food over the high-fat food. Moreover, THC significantly increased high-carbohydrate food feeding bout duration to specifically increase time spent eating the high-carbohydrate food. Together, this provides evidence for THC altering preference for specific macronutrients. Alternatively, in the satiated state THC may increase preference for the less energy dense, less satiating high-carbohydrate food so they can eat more of it to fulfill the need to eat driven by the motivational effects of THC.

Overall, we show that THC vapour exposure alters macronutrient specific food preferences, but the direction in which preference is altered may be dependent on the state of the animal and associated baseline macronutrient preference. Furthermore, in both food choice experiments, THC vapour altered preference in such a way that it abolished pre-existing food preferences so that animals no longer preferred one food over the other. It’s also possible that THC vapour alters palatability and/or food reward value so animals can no longer discriminate between the foods.

Following THC vapour inhalation, we consistently observed that females had significantly higher THC and metabolite concentrations in the blood and brain compared to males. These differences are significant even when concentrations are normalised to body weight, suggesting that this is not solely a consequence of females being smaller in size. Females having higher THC concentrations has been reported in rodent studies injecting THC [49, 50] and is speculated to be a result of THC being sequestered into fat to a higher proportion in males [50]. However, this finding is not consistent between studies. The sex differences we see in THC metabolite concentrations are however consistent with both rodent and human studies [35, 49–52], and imply that females metabolize THC at a faster rate than males, likely due to a sex difference in liver metabolic enzymes [53].

Most cannabis feeding studies exclusively use male subjects, therefore sex differences in cannabis-induced feeding behaviours remain relatively unexplored. For most of this study, we showed that there were no obvious sex differences in the ability of THC to alter food intake, which was surprising as THC has dose dependent effects on feeding behaviours [10, 19]. However, we do see a sex difference with THC effects on macronutrient preferences in standard conditions, where females show a stronger preference for the high-fat food compared to males. This sex difference can be explained by females having higher blood and brain THC concentrations to males. Subtle sex differences in feeding patterns were also noted, for example, THC increases the number of high-carbohydrate food feeding bouts in males more than females, but these differences were not consistent throughout experiments. Irrespective of treatment, certain expected sex differences were detected, such as total amount of food eaten or body weight change.

Furthermore, we did not measure the effects of THC on feeding behaviours during each estrus cycle stage, however, feeding effects were consistent between vapour exposure days and previous studies have not detected THC vapour-induced behavioural differences between estrus and diestrus phases [54].

Ultimately the mechanism underlying THC-driven feeding remains unknown. THC induces its physiological effects through the endocannabinoid system, a modulatory signalling system comprised of receptors, enzymes, and ligands [55], that is involved in numerous homeostatic processes, including energy balance [56, 57]. THC is a partial CB1R agonist, thereby universal CB1R antagonism blocks the appetitive effects of THC [22, 58, 59]. CB1R are expressed across the brain [60, 61] as well as in certain peripheral tissues [62]. THC can cross the blood brain barrier, so THC has the potential to act through peripheral and/or central CB1R to induce its feeding effects.

From a peripheral standpoint, THC could alter levels of circulating appetite and metabolism associated hormones through CB1R expressed within various endocrine organs, including the gastrointestinal system, adipose tissue, and pancreas [63]. One potential mechanism being that THC may promote secretion of the hunger hormone ghrelin from the stomach [64–67]. Additionally, THC could suppress secretion of satiety hormones, however, evidence for cannabis altering energy balance hormones is not consistent between studies [9, 24, 66, 68].

For potential central mechanisms, THC may alter the activity of energy balance and reward regulating brain circuits. CB1R are expressed in the hypothalamic arcuate nucleus (ARC) [69], a major hub for appetite regulation [70]. Specifically within the ARC, CB1R are expressed on GABAergic inputs to appetite stimulating AgRP neurons [71], as well as on appetite inhibiting POMC neurons [72]. Evidence suggests that CB1R agonists can stimulate feeding behaviours through these neuronal populations [69, 71, 72], however, it is yet to be elucidated if THC also acts through these circuits. Furthermore, CB1R are densely expressed in the nucleus accumbens shell (NAcSh) [61], a region heavily involved in reward and food motivation [73]. Like the effects of THC, endocannabinoid activation of NAcSh CB1R stimulates food intake [74] and increases hedonic responses to sucrose [75]. Thereby, THC could also act through a NAcSh pathway to influence feeding behaviours. Ultimately, additional research is required to investigate these potential peripheral and central mechanisms.

For decades, cannabis has had putative therapeutic potential for stimulating appetite. Yet currently the literature is mixed regarding the effects of cannabis use on total energy intake and body weight. In rodent models, inhaled THC can increase total daily energy intake [18, 29, 76], but have no impact on bodyweight [18, 77]. Furthermore, in rodent models of activity-based anorexia, injected THC can acutely increase total energy intake and reduce body weight loss [78–80]. Currently in the clinic, there is variable evidence that cannabis or synthetic cannabinoids can stimulate appetite in cachectic cancer or anorexia nervosa patients [81, 82]. However, around 20% of North American medical cannabis users self-report using cannabis to stimulate appetite [7], higher rates of cannabis use are associated with higher daily caloric intake [83] and cannabis-associated weight gain has been reported in cachectic patients [84–86].

Conversely, chronically administered THC is associated with reduced body weight gain in rodents [31, 87], and regular cannabis users are reported to have lower incidences of obesity compared to the general population [30, 88]. Specifically, regular cannabis use is associated with lower body mass index, waist circumference and fat mass, as well as improved metabolic profiles, including lower circulating insulin levels and reduced insulin resistance scores [89, 90]. Furthermore, in rodents, inhaled THC at higher doses can decrease food intake [19] and increase energy expenditure [18], and chronic injected THC has protective effects on diet-induced obesity development [91]. In alignment with the latter thread of evidence, we show that in our model, once daily THC vapour inhalation acutely increases food intake, which is compensated for to maintain total daily energy intake, and in some cases, is overcompensated for, decreasing total energy intake.

The effects of cannabis use on metabolic health and obesity risks are speculated to be dependent on several factors including differing patterns of cannabis use, specifically with regards to THC dose, the duration and daily frequency of use, and the length of use (i.e., acute vs chronic) [34]. This highlights the necessity for conducting more appropriately designed, controlled, and highly powered pre-clinical and clinical studies assessing the effects of cannabis use regimens on energy balance in the short and long-term, as well as its potential for alleviating cachexia in the clinic. Our food preference data also implies that THC vapour inhalation may not acutely alter energy intake, but it can alter the energy source. Thereby, additional research is needed to assess whether cannabis causes maladaptive eating patterns by driving the consumption of more palatable, energy dense, ‘unhealthy’ foods.

In summary, we have characterised the effects of THC vapour inhalation on rat feeding behaviours, including macronutrient-specific food preferences. This research is important for assessing the energy balance associated consequences of cannabis use, knowledge which is beneficial for both medial and recreational cannabis users. Further investigation is necessary to uncover the association between cannabis use regimens and energy-balance effects, and to decipher the mechanisms underlying cannabis-driven feeding behaviour alterations.

## Acknowledgements

The authors thank La Jolla Alcohol Research Group for their assistance with the vapour exposure system and the University of Calgary Southern Alberta Mass Spectrometry Facility for running samples through mass spectrometry. The authors also thank all who provided support with these experiments, including other Hill lab members and the University of Calgary animal care staff.

## Author Contributions

C.H. and M.N.H. formulated the project and designed the experiments. C.H. undertook all vapour exposure, behavioural experiments, and sample collection. S.L.B. helped with the vapour exposure protocol, and prepared samples and analyzed data for mass spectrometry. C.H., L.J., V.M. and J.B.B. scored feeding behaviour videos. C.H. analyzed all behavioural data and wrote the manuscript. All authors were involved in revising the manuscript.

---

**Supplementary Figure 1.**
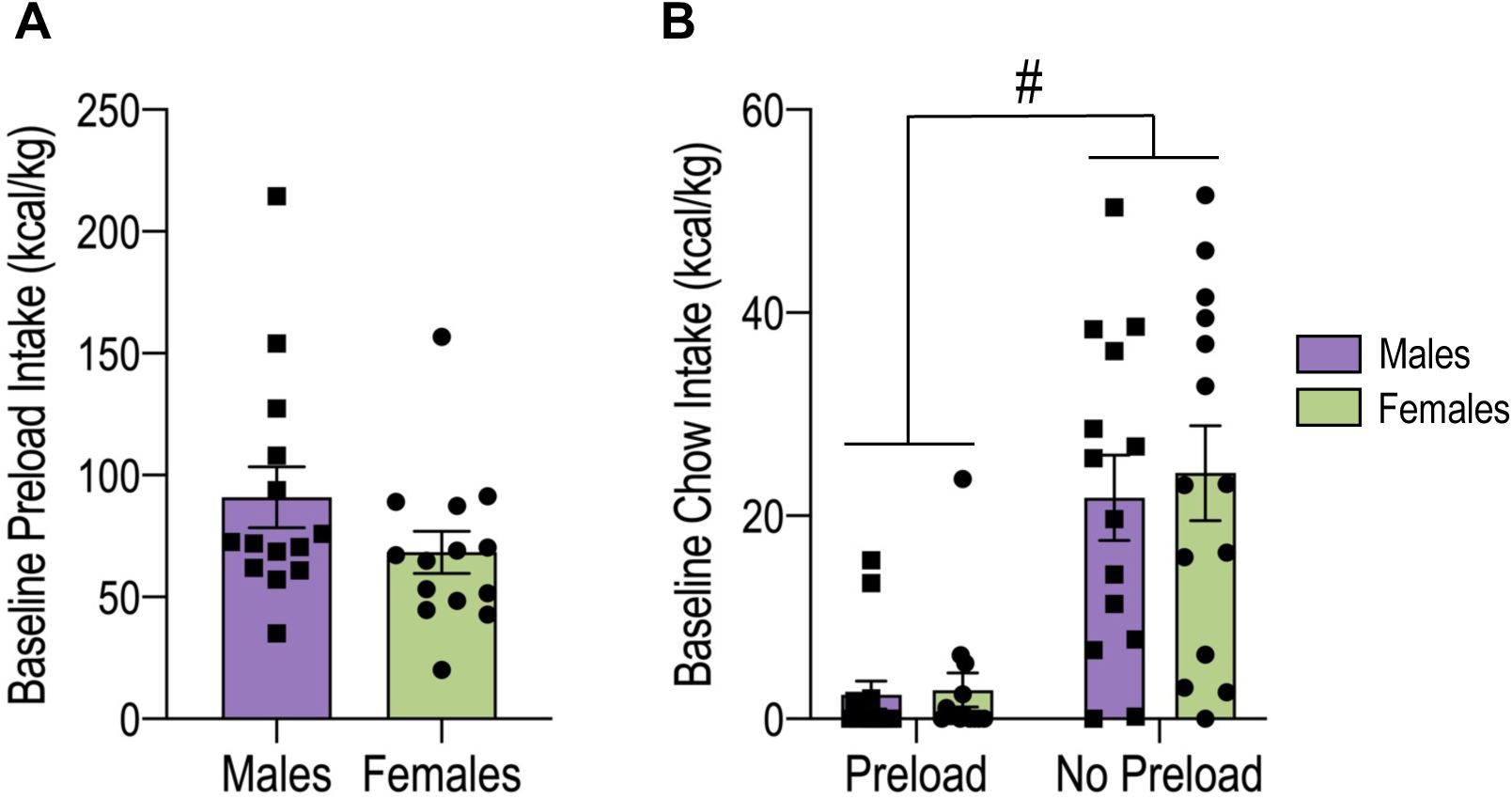
Preload consumption suppresses subsequent chow intake. **(A)** Preload intake at baseline (day 5; rats not exposed to vapour) for male and female rats (unpaired two-tailed t-test, p=0.15). **(B)** 60-minute baseline (no vapour exposure) chow intake for male and female rats with two-hour access to preload or no access to preload (2-way ANOVA, effect of preload: F(1,52)=37.68, p<0.001 (# no preload > preload)). No preload group data from the initial chow feeding experiment outlined in Figure 1. Data normalized to body weight and shown as mean ± SEM. N=14 in each group. # effect of preload.

**Supplementary Figure 2.**
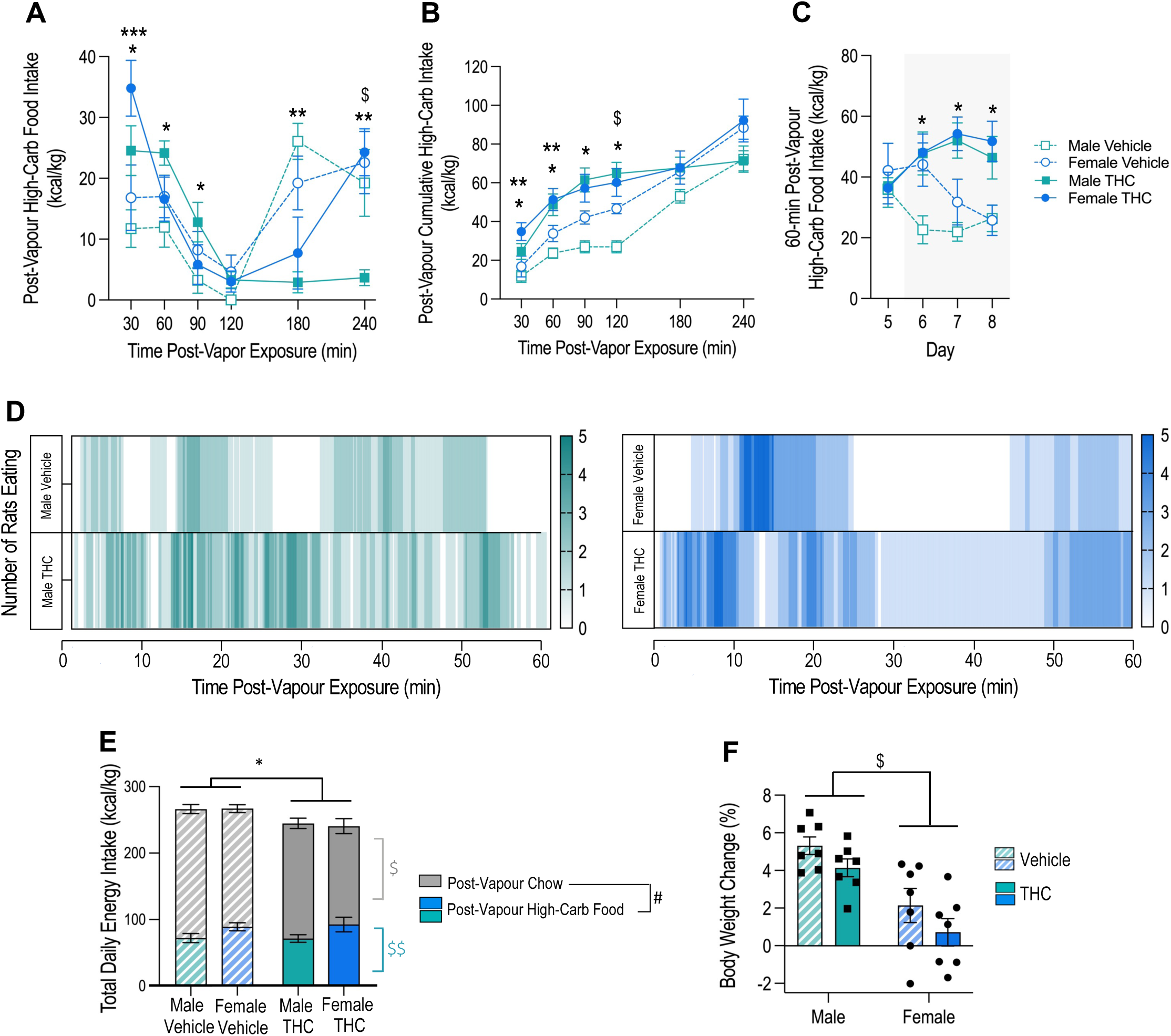
THC vapour acutely increases high-carbohydrate food intake. **(A)** Day 4-6 mean post-vapour high-carbohydrate food intake over time for THC and vehicle groups (RM 3-way ANOVA; interaction effect of time, sex and treatment: F(3.33,79.86)=2.95, p=0.033 (post-hoc: * male THC > male vehicle at t=30, 60 & 90-minutes, ** male vehicle > male THC at t=180 & 240-minutes, *** female THC > female vehicle at t=30-minutes, $ female THC > male THC at t=240-minutes, p<0.05)). **(B)** Day 4-6 mean post-vapour cumulative high-carbohydrate food intake for THC and vehicle groups (RM 3-way ANOVA; interaction effect of time, sex and treatment: F(2.87,68.96)=3.94, p=0.013 (post-hoc: * male THC > male vehicle at t=30, 60, 90 & 120-minutes, ** female THC > female vehicle at t=30 & 60-minutes, $ female vehicle > male vehicle at t=120-minutes, p<0.05)). **(C)** 60-minute high-carbohydrate food intake for day 5 (baseline, no vapour exposure) and 6-8 (post-vapour exposure; highlighted in grey) for THC and vehicle groups (RM 3-way ANOVA, interaction effect of time and treatment: F (3,72)=6.65, p<0.001 (post-hoc: * THC > vehicle on day 6-8, p<0.05)). **(D)** Representative heat maps showing the number and pattern of rats eating high-carbohydrate food within the first 60-minutes after vapour exposure. **(E)** Day 6-8 mean 24-hour energy intake for rats given post-vapour access to high-carbohydrate food (RM 3-way ANOVA; *total intake (high-carbohydrate food + chow) -* effect of treatment: F(1,24)=11.01, p=0.003 (* Vehicle > THC); *total intake of each food -* interaction effect of food and sex: F(1,24)=7.90, p=0.01 (post-hoc: $ male chow intake > female chow intake, $$ female high-carb food intake > male high-carb food intake, # male & female chow intake > male & female high-carb food intake, p<0.05)). **(F)** Body weight change (%) across vapour exposure days (day 6-8) (2-way ANOVA; effect of sex: F(1,24)=24.30, p<0.001 ($ male > female)). All high-carbohydrate food intake data normalized to body weight and shown as mean ± SEM. N=7 in each group. * effect of treatment, $ effect of sex, # effect of food type.

**Supplementary Figure 3.**
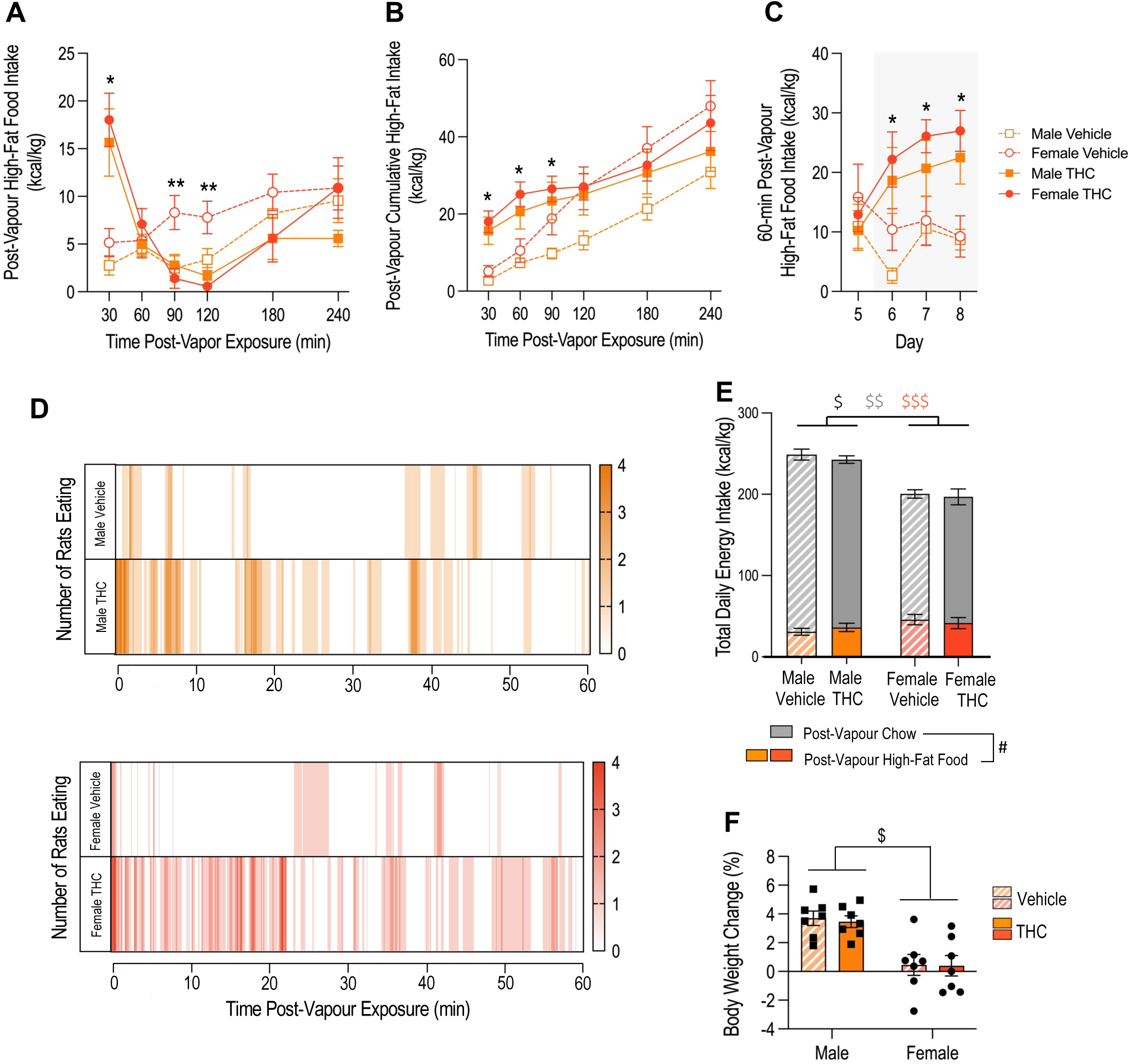
THC vapour acutely increases high-fat food intake. **(A)** Day 4-6 mean post-vapour high-fat food intake over time for THC and vehicle groups (RM 3-way ANOVA; interaction effect of time and treatment: F(3.67,87.96)=13.87, p<0.001 (post-hoc: * THC > vehicle at t=30-minutes, ** vehicle > THC at t=90 & 120-minutes, p<0.05). **(B)** Day 4-6 mean post-vapour cumulative high-fat food intake for THC and vehicle groups (RM 3-way ANOVA; interaction effect of time and treatment: F(1.86,44.59)=8.00, p=0.001 (post-hoc: * THC > vehicle at t=30, 60 & 90-minutes, p<0.05). **(C)** Mean 60-minute high-fat food intake for day 5 (baseline, no vapour exposure) and 6-8 (post-vapour exposure; highlighted in grey) for THC and vehicle groups (RM 3-way ANOVA; interaction effect of day and treatment: F(3,72)=9.56, p<0.001 (post-hoc: * THC > vehicle on days 6-8, p<0.05). **(D)** Representative heat maps showing the number and pattern of rats eating high-fat food within the first 60-minutes after vapour exposure. **(E)** Day 6-8 mean 24-hour energy intake for rats given post-vapour access to high-fat food (RM 3-way ANOVA; *total intake (high-fat food + chow)* - effect of sex: F(1,24)=59.1, p<0.001 ($ male > female); *total intake of each food* - interaction effect of food and sex: F(1,24)=28.04, p<0.001 (post-hoc: $$ male chow intake > female chow intake, $$$ female high-fat food intake > male high-fat food intake, # male & female chow intake > male & female high-fat food intake, p<0.05)). **(F)** Body weight change (%) across vapour exposure days (day 6-8) (2-way ANOVA; effect of sex: F(1,24)=27.42, p<0.001 ($ male > female)). All high-fat food intake data normalized to body weight and shown as mean ± SEM. N=7 in each group. * effect of treatment, $ effect of sex, # effect of food type.

**Supplementary Figure 4.**
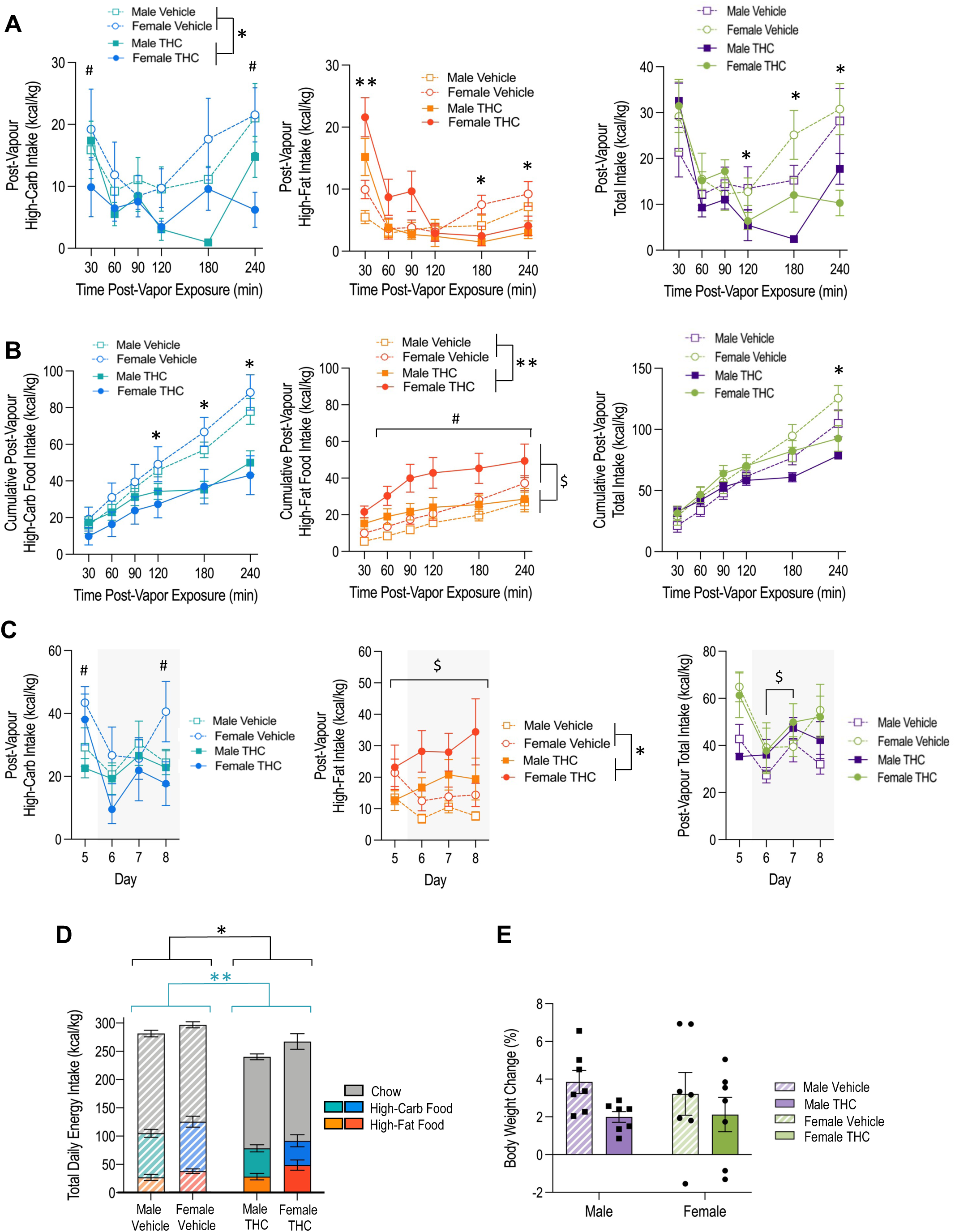
THC acutely and selectively increases high-fat food intake when given a choice between a high-fat and a high-carbohydrate food. **(A)** Day 6-8 mean post-vapour intake of high-carbohydrate food, high-fat food, and total food (high-carbohydrate food + high-fat food) intake over time for THC and vehicle groups (RM 3-way ANOVA; *high-carbohydrate –* effect of vapour treatment: F(1,24)=17.98, p<0.001 (* vehicle > THC) & effect of time: F(1,120)=4.50, p<0.001 (post-hoc: # t=30 & 240-min > t=120-min, p<0.05); *high-fat –* interaction effect of time and vapour treatment: F(1,88.94)=12.25, p<0.001 (post-hoc: ** THC > vehicle at t=30, * vehicle > THC at t=180 & 240-min, p<0.05); *total* – interaction effect of time and vapour treatment: F(5,120)=3.94, p=0.002 (post-hoc: * vehicle > THC at t=120, 180 & 240-min, p<0.05). (B) Day 6-8 mean cumulative post-vapour intake of high-carbohydrate food, high-fat food, and total food (high-carbohydrate food + high-fat food) intake over time for THC and vehicle groups (RM 3-way ANOVA; *high-carbohydrate –* interaction effect of time and treatment: F(2.77,66.35)=14.27, p<0.001 (* vehicle > THC at t=120, 180 & 240-min, p<0.05); *high-fat* – effect of time: F(2.06,41.03)=57.95, p<0.001 (# t=30 < t=60, 90, 120, 180 & 240-min, p<0.001) & effect of treatment: F(1,24)=7.84, p=0.01 (** THC > vehicle) & effect of sex: F(1,24)=6.36, p=0.019 ($ female > male); *total* – interaction effect of time and vapour treatment: F(2.60,64.31)=16.44, p<0.001 (post-hoc: * vehicle > THC at t=240-min, p<0.05). (C) Mean 60-minute post-vapour intake of high-carbohydrate food, high-fat food, and total food (high-carbohydrate food + high-fat food) for day 5 (baseline: no vapour exposure) and 6-8 (post-vapour exposure; highlighted in grey) for THC and vehicle groups (RM 3-way ANOVA; *high-carbohydrate –* effect of time F(1,45.04)=4.48, p=0.019 (post-hoc: # day 5 & 8 > day 6, p<0.05); *high-fat –* effect of vapour treatment: F(1,24)=10.09, p=0.004 (* THC > vehicle) & effect of sex: F(1,24)=6.74, p=0.016 ($ female > male); *total* – effect of day: F(2.21,52.99)=4.29, p=0.016 (post-hoc: $ day 3 > day 2, p<0.05). (D) Day 6-8 mean 24-hour energy intake for THC and vehicle groups (RM 3-way ANOVA; *total intake (high-fat food + high-carbohydrate food + chow) –* effect of treatment: F(1,24)=10.43, p=0.004 (* vehicle > THC) & effect of sex: F(1,24)=4.45, p=0.046 ($ female > male); *total intake of each food –* interaction effect of vapour treatment and food type: F(1,48)=6.82, p=0.002 (post-hoc: ** THC high-carbohydrate food intake < vehicle high-carbohydrate food intake, p<0.01)). (E) Body weight change (%) across vapour exposure days (day 6-8) (2-way ANOVA; no significant effect of treatment: F(1,24)=3.4, p=0.077). Food intake data normalized to body weight and shown as mean ± SEM. N=7 in each group. # effect of time, * effect of treatment, $ effect of sex

**Supplementary Figure 5.**
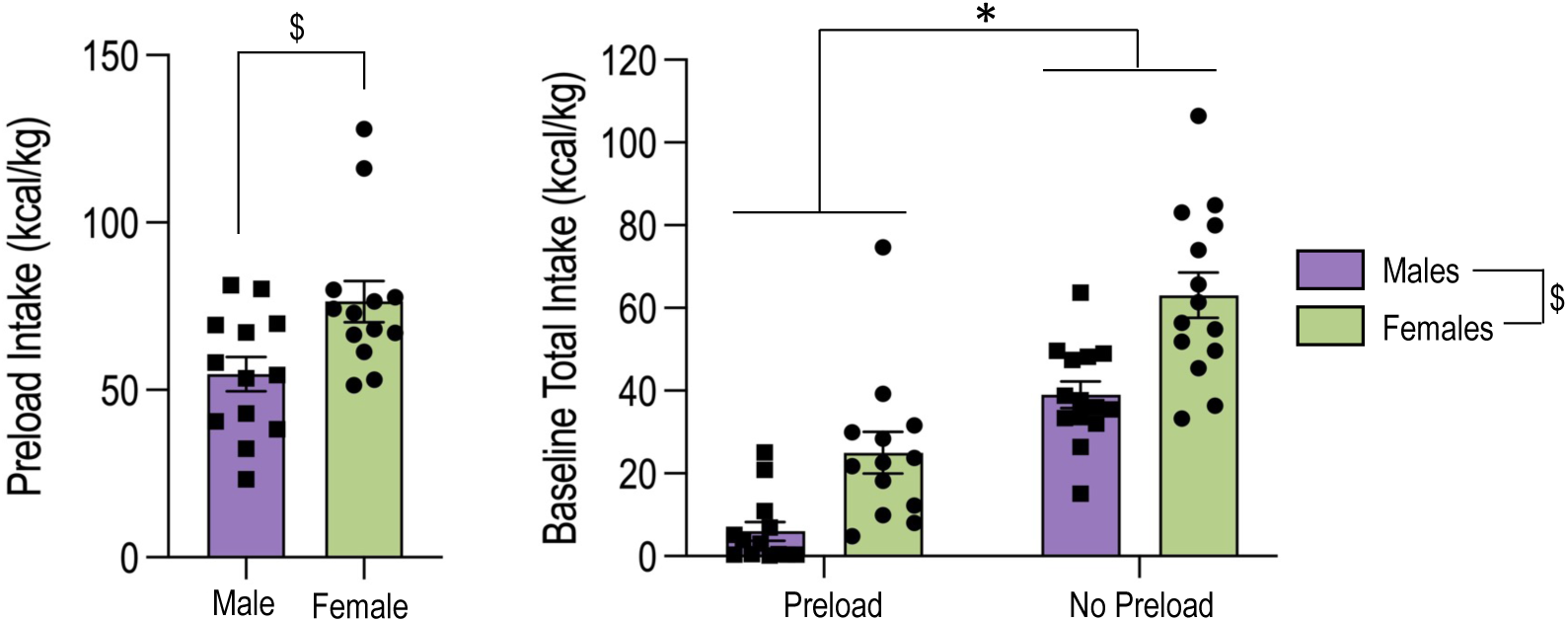
Preload consumption suppresses subsequent high-carbohydrate and high-fat food intake. **(A)** Preload intake at baseline (day 5; rats not exposed to vapour) for male and female rats (unpaired two-tailed t-test; $ p=0.012). **(B)** 60-minute baseline (no vapour exposure) total intake (high-carbohydrate food + high-fat food) for male and female rats with two-hour access to preload or no access to preload (2-way ANOVA; effect of preload: F(1,50)=70.65, p<0.001 (* no preload > preload) & effect of sex: F(1,50)=25.91, p<0.001 ($ females > males)). No preload group data from the initial food preference experiment outlined in Figure 5. Data normalized to body weight and shown as mean ± SEM. N=14 in the no preload groups and n=13 in the preload groups; one rat in both the male and female preload groups were excluded based on meeting outlier criteria (>2.5 SD away from the mean).

**Supplementary Figure 6.**
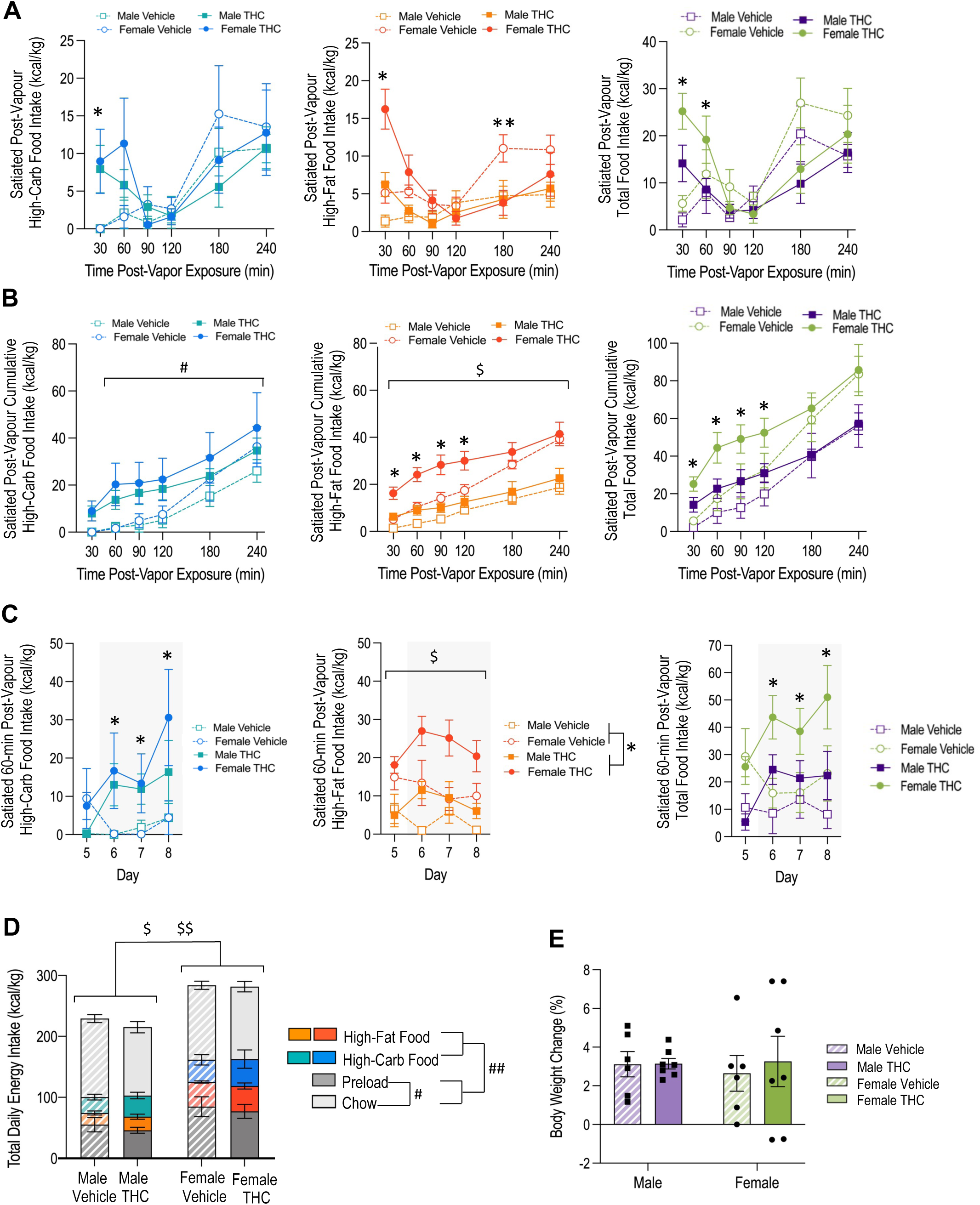
In the satiated state, THC acutely increases intake of both high-fat and high-carbohydrate foods when given a choice between both. **(A)** Day 6-8 mean post-vapour intake of high-carbohydrate food, high-fat food, and total food (high-carbohydrate food + high-fat food) intake over time for satiated THC and vehicle groups (RM 3-way ANOVA; *high-carb –* interaction effect of time and treatment: F(3.2,96.88)=2.80, p=0.043 (post-hoc: * THC > vehicle at t=30-min, p<0.01); *high-fat –* interaction effect of time and treatment: F(5,110)=7.13, p<0.001 (post-hoc: * THC > vehicle at t=30-min, ** vehicle > THC at t=180-min, p<0.05); *total* - interaction effect of time and treatment: F(2.53,55.62)=6.72, p=0.001 (post-hoc: * THC > vehicle at t=30 & 60-min, p<0.05). (B) Day 6-8 mean cumulative post-vapour intake of high-carbohydrate food, high-fat food, and total food (high-carbohydrate food + high-fat food) intake over time for satiated THC and vehicle groups (RM 3-way ANOVA; *high-carbohydrate –* effect of time : F(1.93,42.41)=44.94, p<0.001 (# t=30 < t=60, 90, 120, 180 & 240-min, p<0.05); *high-fat –* interaction effect between time and treatment: F(2.17,47.84)=3.62, p=0.031 (* THC > vehicle at t=30, 60, 90 & 120-minutes, p<0.05) & interaction effect between time and sex: F(2.17,47.84)=5.98, p=0.004 ($ female > male at all time points, p<0.05); *total* – interaction effect of time and treatment: F(2.25,49.45)=3.23, p=0.043 (post-hoc: * THC > vehicle at t=30, 60, 90 & 120-min, p<0.05). (C) Mean 60-minute post-vapour intake of high-carbohydrate food, high-fat food and total food (high-carbohydrate food + high-fat food) for day 5 (baseline: no vapour exposure) and 6-8 (post-vapour exposure; highlighted in grey) for satiated THC and vehicle groups (RM 3-way ANOVA; *high-carb –* interaction effect of time and treatment: F(2.65,58.33)=3.04, p=0.042 (post-hoc: * THC > vehicle on days 6-8, p<0.01); *high-fat –* effect of treatment: F(1,22)=13.12, p=0.022 (* THC > vehicle) & effect of sex: F(1,22)=30.05, p<0.001 ($ female > male); *total* - interaction effect of time and treatment: F(3,66)=5.24, p=0.003 (post-hoc: * THC > vehicle on days 6-8, p<0.05). (D) Day 6-8 mean 24-hour energy intake for THC and vehicle groups (RM 3-way ANOVA; *total intake (preload + high-fat food + high-carbohydrate food + chow) –* effect of sex: F(1,22)=30.98, p<0.001 ($ female > male); *total intake of each food –* effect of sex: F(1,22)=30.98, p<0.001 ($$ female > male) & effect of food: F(2.16,47.64)=73.67, p<0.001 (post-hoc: # chow > preload, ## chow & preload > high-carbohydrate & high fat food, p<0.05)). (E) Body weight change (%) across vapour exposure days (day 6-8) (2-way ANOVA; no significant effect of treatment: F(1,22)=0.13, p=0.72). Food intake data normalized to body weight and shown as mean ± SEM. N=7 in THC groups and n=6 in vehicle groups; one rat in both the male and female vehicle group were excluded based on meeting outlier criteria (>2.5 SD away from the mean). # effect of food/time, * effect of treatment, $ effect of sex.

**Supplementary Table 1.**
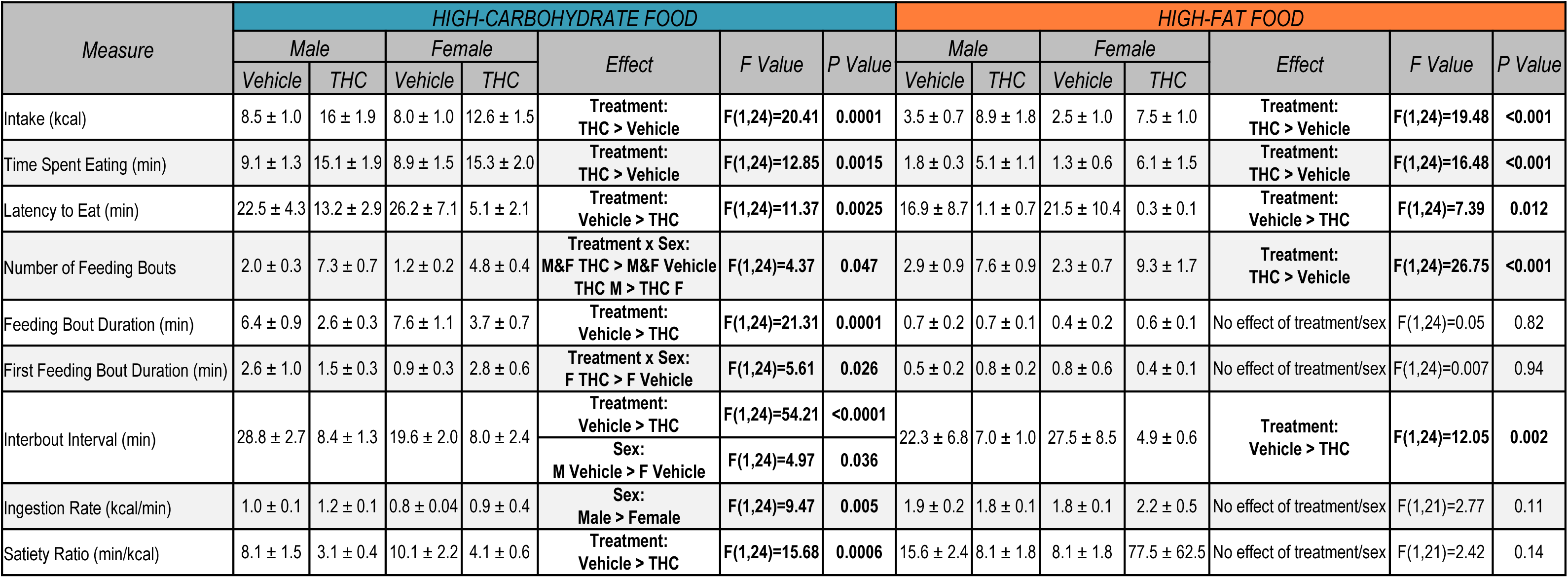
THC increases high-carbohydrate or high-fat food intake by increasing latency to eat and feeding bout frequency to increase overall time spent eating. Day 4-6 mean feeding behaviour parameters measured within the first 60-minutes after vapour exposure. Data shown as mean ± SEM and analysed using a two-way ANOVA (for parameters with no effect, only the F and P values for treatment comparison are shown). N=7 in each group, except from ingestion rate and satiety ratio in the high-fat food access experiment (male vehicle n=6, female vehicle n=5, male & female THC n=7) - these parameters can’t be calculated from animals that don’t eat. Feeding bouts: arbitrarily defined periods of eating separated by 30-seconds or more. Ingestion rate: high-carbohydrate food calories consumed divided by time spent eating. Satiety ratio: time spent not eating divided by high-carbohydrate food calories consumed. M: male. F: female.

## References

1. United Nations Office on Drugs and Crime., Drug market trends of Cannabis and Opioids. World Drug Report 2022; https://www.unodc.org/res/wdr2022/MS/WDR22_Booklet_3.pdf.

2. Memedovich, K.A., et al., The adverse health effects and harms related to marijuana use: an overview review. Canadian Medical Association Open Access Journal, 2018. 6(3): p. E339–E346.

3. Goodman, S., et al., Prevalence and forms of cannabis use in legal vs. illegal recreational cannabis markets. International Journal of Drug Policy, 2020. 76: p. 102658.

4. Manthey, J., et al., Public health monitoring of cannabis use in Europe: prevalence of use, cannabis potency, and treatment rates. The Lancet Regional Health-Europe, 2021. 10: p. 100227.

5. Government of Canada., Canadian Cannabis Survey 2021; https://publications.gc.ca/collections/collection_2021/sc-hc/H21-312-2021-3-eng.pdf.

6. ElSohly, M.A., et al., A comprehensive review of cannabis potency in the United States in the last decade. Biological Psychiatry: Cognitive Neuroscience and Neuroimaging, 2021. 6(6): p. 603–606.

7. Leung, J., et al., Prevalence and self-reported reasons of cannabis use for medical purposes in USA and Canada. Psychopharmacology, 2022. 239(5): p. 1509–1519.

8. Legare, C.A., W.M. Raup-Konsavage, and K.E. Vrana, Therapeutic potential of cannabis, cannabidiol, and cannabinoid-based pharmaceuticals. Pharmacology, 2022: p. 1–19.

9. Weltens, N., et al., Effect of acute Δ9-tetrahydrocannabinol administration on subjective and metabolic hormone responses to food stimuli and food intake in healthy humans: a randomized, placebo-controlled study. The American Journal of Clinical Nutrition, 2019. 109(4): p. 1051–1063.

10. Williams, C.M., P.J. Rogers, and T.C. Kirkham, Hyperphagia in pre-fed rats following oral Δ9-THC. Physiology & behavior, 1998. 65(2): p. 343–346.

11. Brisbois, T., et al., Delta-9-tetrahydrocannabinol may palliate altered chemosensory perception in cancer patients: results of a randomized, double-blind, placebo-controlled pilot trial. Annals of Oncology, 2011. 22(9): p. 2086–2093.

12. De Bruijn, S.E., et al., Explorative placebo-controlled double-blind intervention study with low doses of inhaled Δ9-tetrahydrocannabinol and cannabidiol reveals no effect on sweet taste intensity perception and liking in humans. Cannabis and cannabinoid research, 2017. 2(1): p. 114–122.

13. Dovey, T.M., et al., Alterations in taste perception due to recreational drug use are due to smoking a substance rather than ingesting it. Appetite, 2016. 107: p. 1–8.

14. Foltin, R.W., M.W. Fischman, and M.F. Byrne, Effects of smoked marijuana on food intake and body weight of humans living in a residential laboratory. Appetite, 1988. 11(1): p. 1–14.

15. Higgs, S., et al., Differential effects of two cannabinoid receptor agonists on progressive ratio responding for food and free-feeding in rats. Behavioural pharmacology, 2005. 16(5-6): p. 389–393.

16. Moore, C.F., et al., Appetitive, antinociceptive, and hypothermic effects of vaped and injected Δ-9-tetrahydrocannabinol (THC) in rats: exposure and dose-effect comparisons by strain and sex. Pharmacology Biochemistry and Behavior, 2021. 202: p. 173116.

17. Solinas, M. and S.R. Goldberg, Motivational effects of cannabinoids and opioids on food reinforcement depend on simultaneous activation of cannabinoid and opioid systems. Neuropsychopharmacology, 2005. 30(11): p. 2035–2045.

18. Freels, T.G., et al., Vaporized cannabis extracts have reinforcing properties and support conditioned drug-seeking behavior in rats. Journal of Neuroscience, 2020. 40(9): p. 1897–1908.

19. Manwell, L.A., et al., A vapourized δ9-tetrahydrocannabinol (δ9-THC) delivery system part II: Comparison of behavioural effects of pulmonary versus parenteral cannabinoid exposure in rodents. Journal of pharmacological and toxicological methods, 2014. 70(1): p. 112–119.

20. Koch, J.E., Δ9-THC stimulates food intake in Lewis rats: effects on chow, high-fat and sweet high-fat diets. Pharmacology Biochemistry and Behavior, 2001. 68(3): p. 539–543.

21. De Luca, M.A., et al., Cannabinoid facilitation of behavioral and biochemical hedonic taste responses. Neuropharmacology, 2012. 63(1): p. 161–168.

22. Jarrett, M.M., J. Scantlebury, and L.A. Parker, Effect of Δ9-tetrahydrocannabinol on quinine palatability and AM251 on sucrose and quinine palatability using the taste reactivity test. Physiology & behavior, 2007. 90(2-3): p. 425–430.

23. Brutman, J.N., et al., Vapor cannabis exposure promotes genetic plasticity in the rat hypothalamus. Scientific Reports, 2019. 9(1): p. 1–12.

24. Muniyappa, R., et al., Metabolic effects of chronic cannabis smoking. Diabetes care, 2013. 36(8): p. 2415–2422.

25. Farrimond, J.A., et al., Cannabis constituents modulateΔ 9-tetrahydrocannabinol-induced hyperphagia in rats. Psychopharmacology, 2010. 210(1): p. 97–106.

26. Farrimond, J.A., B.J. Whalley, and C.M. Williams, A low-Δ9tetrahydrocannabinol cannabis extract induces hyperphagia in rats. Behavioural Pharmacology, 2010. 21(8): p. 769–772.

27. Bellocchio, L., et al., Bimodal control of stimulated food intake by the endocannabinoid system. Nature neuroscience, 2010. 13(3): p. 281–283.

28. Moore, C.F., et al., Translational models of cannabinoid vapor exposure in laboratory animals. Behavioural Pharmacology, 2022. 33(2-3): p. 63–89.

29. Foltin, R.W., J.V. Brady, and M.W. Fischman, Behavioral analysis of marijuana effects on food intake in humans. Pharmacology Biochemistry and Behavior, 1986. 25(3): p. 577–582.

30. Hayatbakhsh, M.R., et al., Cannabis use and obesity and young adults. The American journal of drug and alcohol abuse, 2010. 36(6): p. 350–356.

31. Manning, F.J., et al., Inhibition of normal growth by chronic administration of Δ9-tetrahydrocannabinol. Science, 1971. 174(4007): p. 424-426.

32. Badowski, M.E. and P.K. Yanful, Dronabinol oral solution in the management of anorexia and weight loss in AIDS and cancer. Therapeutics and clinical risk management, 2018. 14: p. 643.

33. McKee, K.A., et al., Potential therapeutic benefits of cannabinoid products in adult psychiatric disorders: A systematic review and meta-analysis of randomised controlled trials. Journal of Psychiatric Research, 2021. 140: p. 267–281.

34. Fearby, N., S. Penman, and P. Thanos, Effects of Δ9-Tetrahydrocannibinol (THC) on Obesity at Different Stages of Life: A Literature Review. International Journal of Environmental Research and Public Health, 2022. 19(6): p. 3174.

35. Baglot, S.L., et al., Pharmacokinetics and central accumulation of delta-9-tetrahydrocannabinol (THC) and its bioactive metabolites are influenced by route of administration and sex in rats. Scientific Reports, 2021. 11(1): p. 1–14.

36. McLaughlin, R.J., Toward a translationally relevant preclinical model of cannabis use. Neuropsychopharmacology, 2018. 43(1): p. 213.

37. Farrimond, J.A., B.J. Whalley, and C.M. Williams, Non-Δ9tetrahydrocannabinol phytocannabinoids stimulate feeding in rats. Behavioural Pharmacology, 2012. 23(1): p. 113–117.

38. Baglot, S.L., et al., Maternal-fetal transmission of delta-9-tetrahydrocannabinol (THC) and its metabolites following inhalation and injection exposure during pregnancy in rats. Journal of Neuroscience Research, 2022. 100(3): p. 713–730.

39. Lindgren, J.-E., et al., Clinical effects and plasma levels of Δ9-tetrahydrocannabinol (Δ9-THC) in heavy and light users of cannabis. Psychopharmacology, 1981. 74(3): p. 208–212.

40. Goebel, M., et al., Central nesfatin-1 reduces the nocturnal food intake in mice by reducing meal size and increasing inter-meal intervals. Peptides, 2011. 32(1): p. 36–43.

41. Zheng, H., et al., Meal patterns, satiety, and food choice in a rat model of Roux-en-Y gastric bypass surgery. American Journal of Physiology-Regulatory, Integrative and Comparative Physiology, 2009. 297(5): p. R1273–R1282.

42. Zorrilla, E.P., et al., Measuring meals: structure of prandial food and water intake of rats. American Journal of Physiology-Regulatory, Integrative and Comparative Physiology, 2005. 288(6): p. R1450–R1467.

43. Low, A.Y., et al., Reverse-translational identification of a cerebellar satiation network. Nature, 2021. 600(7888): p. 269-273.

44. Williams, C.M. and T.C. Kirkham, Observational analysis of feeding induced by Δ9-THC and anandamide. Physiology & behavior, 2002. 76(2): p. 241–250.

45. Escartín-Pérez, R.E., et al., Role of cannabinoid CB1 receptors on macronutrient selection and satiety in rats. Physiology & behavior, 2009. 96(4-5): p. 646–650.

46. Radziszewska, E., M. Wolak, and E. Bojanowska, Concurrent pharmacological modification of cannabinoid-1 and glucagon-like peptide-1 receptor activity affects feeding behavior and body weight in rats fed a free-choice, high-carbohydrate diet. Behavioural Pharmacology, 2014. 25(1): p. 53–60.

47. Wierucka-Rybak, M., M. Wolak, and E. Bojanowska, The effects of leptin in combination with a cannabinoid receptor 1 antagonist, AM 251, or cannabidiol on food intake and body weight in rats fed a high-fat or a free-choice high sugar diet. J Physiol Pharmacol, 2014. 65(4): p. 487-496.

48. Higuchi, S., et al., Increment of hypothalamic 2-arachidonoylglycerol induces the preference for a high-fat diet via activation of cannabinoid 1 receptors. Behavioural brain research, 2011. 216(1): p. 477–480.

49. Ruiz, C.M., et al., Pharmacokinetic, behavioral, and brain activity effects of Δ9-tetrahydrocannabinol in adolescent male and female rats. Neuropsychopharmacology, 2021. 46(5): p. 959–969.

50. Tseng, A.H., J.W. Harding, and R.M. Craft, Pharmacokinetic factors in sex differences in Δ9-tetrahydrocannabinol-induced behavioral effects in rats. Behavioural brain research, 2004. 154(1): p. 77–83.

51. Sholler, D.J., et al., Sex differences in the acute effects of oral and vaporized cannabis among healthy adults. Addiction biology, 2021. 26(4): p. e12968.

52. Wiley, J.L. and J.J. Burston, Sex differences in Δ9-tetrahydrocannabinol metabolism and in vivo pharmacology following acute and repeated dosing in adolescent rats. Neuroscience letters, 2014. 576: p. 51–55.

53. Cooper, Z.D. and R.M. Craft, Sex-dependent effects of cannabis and cannabinoids: a translational perspective. Neuropsychopharmacology, 2018. 43(1): p. 34–51.

54. Javadi-Paydar, M., et al., Effects of Δ9-THC and cannabidiol vapor inhalation in male and female rats. Psychopharmacology, 2018. 235(9): p. 2541–2557.

55. Zou, S. and U. Kumar, Cannabinoid receptors and the endocannabinoid system: signaling and function in the central nervous system. International journal of molecular sciences, 2018. 19(3): p. 833.

56. Aguilera Vasquez, N. and D.E. Nielsen, The Endocannabinoid System and Eating Behaviours: a Review of the Current State of the Evidence. Current Nutrition Reports, 2022: p. 1–10.

57. Ruiz de Azua, I. and B. Lutz, Multiple endocannabinoid-mediated mechanisms in the regulation of energy homeostasis in brain and peripheral tissues. Cellular and molecular life sciences, 2019. 76(7): p. 1341–1363.

58. Järbe, T. and N. DiPatrizio, Δ9-THC induced hyperphagia and tolerance assessment: interactions between the CB1 receptor agonist Δ9-THC and the CB1 receptor antagonist SR-141716 (rimonabant) in rats. Behavioural pharmacology, 2005. 16(5-6): p. 373–380.

59. Williams, C.M. and T.C. Kirkham, Reversal of Δ9-THC hyperphagia by SR141716 and naloxone but not dexfenfluramine. Pharmacology Biochemistry and Behavior, 2002. 71(1-2): p. 333–340.

60. Herkenham, M., et al., Cannabinoid receptor localization in brain. Proceedings of the national Academy of sciences, 1990. 87(5): p. 1932–1936.

61. Tsou, K., et al., Immunohistochemical distribution of cannabinoid CB1 receptors in the rat central nervous system. Neuroscience, 1998. 83(2): p. 393–411.

62. Galiègue, S., et al., Expression of central and peripheral cannabinoid receptors in human immune tissues and leukocyte subpopulations. European journal of biochemistry, 1995. 232(1): p. 54–61.

63. O’Sullivan, S.E., A.S. Yates, and R.K. Porter, The Peripheral Cannabinoid Receptor Type 1 (CB1) as a Molecular Target for Modulating Body Weight in Man. Molecules, 2021. 26(20): p. 6178.

64. Cani, P.D., et al., Potential modulation of plasma ghrelin and glucagon-like peptide-1 by anorexigenic cannabinoid compounds, SR141716A (rimonabant) and oleoylethanolamide. British Journal of Nutrition, 2004. 92(5): p. 757-761.

65. Mazidi, M., et al., The effect of hydroalcoholic extract of Cannabis Sativa on appetite hormone in rat. Journal of Complementary and Integrative Medicine, 2014. 11(4): p. 253–257.

66. Riggs, P.K., et al., A pilot study of the effects of cannabis on appetite hormones in HIV-infected adult men. Brain research, 2012. 1431: p. 46–52.

67. Zbucki, R.L., et al., Cannabinoids enhance gastric X/A-like cells activity. Folia Histochemica et Cytobiologica, 2008. 46(2): p. 219–224.

68. Farokhnia, M., et al., Effects of oral, smoked, and vaporized cannabis on endocrine pathways related to appetite and metabolism: a randomized, double-blind, placebo-controlled, human laboratory study. Translational psychiatry, 2020. 10(1): p. 1–11.

69. Cardinal, P., et al., Hypothalamic CB1 cannabinoid receptors regulate energy balance in mice. Endocrinology, 2012. 153(9): p. 4136–4143.

70. Andermann, M.L. and B.B. Lowell, Toward a wiring diagram understanding of appetite control. Neuron, 2017. 95(4): p. 757–778.

71. Morozov, Y.M., et al., Cannabinoid type 1 receptor-containing axons innervate NPY/AgRP neurons in the mouse arcuate nucleus. Molecular metabolism, 2017. 6(4): p. 374–381.

72. Koch, M., et al., Hypothalamic POMC neurons promote cannabinoid-induced feeding. Nature, 2015. 519(7541): p. 45-50.

73. Morales, I. and K.C. Berridge, ‘Liking’and ‘wanting’in eating and food reward: Brain mechanisms and clinical implications. Physiology & behavior, 2020. 227: p. 113152.

74. Kirkham, T.C., et al., Endocannabinoid levels in rat limbic forebrain and hypothalamus in relation to fasting, feeding and satiation: stimulation of eating by 2-arachidonoyl glycerol. British journal of pharmacology, 2002. 136(4): p. 550–557.

75. Mahler, S.V., K.S. Smith, and K.C. Berridge, Endocannabinoid hedonic hotspot for sensory pleasure: anandamide in nucleus accumbens shell enhances ‘liking’of a sweet reward. Neuropsychopharmacology, 2007. 32(11): p. 2267–2278.

76. Hart, C.L., et al., Comparison of smoked marijuana and oral Δ9-tetrahydrocannabinol in humans. Psychopharmacology, 2002. 164(4): p. 407–415.

77. Nguyen, J.D., et al., Lasting effects of repeated Δ9-tetrahydrocannabinol (THC) vapor inhalation during adolescence in male and female rats. bioRxiv, 2019: p. 426064.

78. Lewis, D.Y. and R.R. Brett, Activity-based anorexia in C57/BL6 mice: Effects of the phytocannabinoid,△ 9-tetrahydrocannabinol (THC) and the anandamide analogue, OMDM-2. European Neuropsychopharmacology, 2010. 20(9): p. 622-631.

79. Scherma, M., et al., Cannabinoid CB1/CB2 receptor agonists attenuate hyperactivity and body weight loss in a rat model of activity-based anorexia. British journal of pharmacology, 2017. 174(16): p. 2682–2695.

80. Verty, A.N., et al., The cannabinoid receptor agonist THC attenuates weight loss in a rodent model of activity-based anorexia. Neuropsychopharmacology, 2011. 36(7): p. 1349–1358.

81. Razmovski-Naumovski, V., et al., Efficacy of medicinal cannabis for appetite-related symptoms in people with cancer: A systematic review. Palliative Medicine, 2022: p. 02692163221083437.

82. Rosager, E.V., C. Møller, and M. Sjögren, Treatment studies with cannabinoids in anorexia nervosa: a systematic review. Eating and Weight Disorders-Studies on Anorexia, Bulimia and Obesity, 2021. 26(2): p. 407–415.

83. Rodondi, N., et al., Marijuana use, diet, body mass index, and cardiovascular risk factors (from the CARDIA study). The American journal of cardiology, 2006. 98(4): p. 478–484.

84. Andries, A., et al., Dronabinol in severe, enduring anorexia nervosa: a randomized controlled trial. International Journal of Eating Disorders, 2014. 47(1): p. 18–23.

85. Bar-Sela, G., et al., The effects of dosage-controlled cannabis capsules on cancer-related cachexia and anorexia syndrome in advanced cancer patients: pilot study. Integrative Cancer Therapies, 2019. 18: p. 1534735419881498.

86. Haney, M., et al., Dronabinol and marijuana in HIV-positive marijuana smokers: caloric intake, mood, and sleep. JAIDS Journal of Acquired Immune Deficiency Syndromes, 2007. 45(5): p. 545–554.

87. Nelson, N.G., et al., CombinedΔ 9-tetrahydrocannabinol and moderate alcohol administration: Effects on ingestive behaviors in adolescent male rats. Psychopharmacology, 2019. 236(2): p. 671–684.

88. Le Strat, Y. and B. Le Foll, Obesity and cannabis use: results from 2 representative national surveys. American journal of epidemiology, 2011. 174(8): p. 929–933.

89. Ngueta, G., et al., Cannabis use in relation to obesity and insulin resistance in the Inuit population. Obesity, 2015. 23(2): p. 290–295.

90. Penner, E.A., H. Buettner, and M.A. Mittleman, The impact of marijuana use on glucose, insulin, and insulin resistance among US adults. The American journal of medicine, 2013. 126(7): p. 583–589.

91. Cluny, N.L., et al., Prevention of diet-induced obesity effects on body weight and gut microbiota in mice treated chronically with Δ9-tetrahydrocannabinol. PloS one, 2015. 10(12): p. e0144270.

